# The cellular landscape of the endochondral bone during the transition to extrauterine life

**DOI:** 10.1101/2023.07.18.549529

**Authors:** Alejandro Díaz Rueda, Irepan Salvador-Martínez, Ismael Sospedra-Arrufat, Ana Alcaina-Caro, Ana Fernández-Miñán, Ana M. Burgos-Ruiz, Ildefonso Cases, Alberto Mohedano, Juan J. Tena, Holger Heyn, Javier Lopez-Rios, Gretel Nusspaumer

## Abstract

The cellular complexity of the endochondral bone underlies its essential and pleiotropic roles during organismal life. While the adult bone has received significant attention, we still lack a deep understanding of the perinatal bone cellulome. Here, we have profiled the full composition of the murine endochondral bone at the single-cell level during the transition from fetal to newborn life and in comparison to the adult tissue, with particular emphasis on the mesenchymal compartment. The perinatal bone contains different fibroblastic clusters with blastema-like characteristics in organizing and supporting skeletogenesis, angiogenesis, and hematopoiesis. Our data also suggests dynamic inter- and intra-compartment interactions as well as a bone marrow milieu that seems prone to anti-inflammation, which we hypothesize is necessary to ensure the proper program of lymphopoiesis and the establishment of central and peripheral tolerance in early life. Our study provides an integrative roadmap for the future design of genetic and cellular functional assays to validate cellular interactions and lineage relationships within the perinatal bone.

## INTRODUCTION

Important changes take place during the transition from intrauterine to extrauterine life. The newborn is reaved of nutrients, heat and the buoyant and pathogen-protected environment of the womb. Among other large organismal changes, lungs are inflated and the cardiovascular flow adjusted as the newborn starts breathing. High hormonal levels are present in preparation for the sharp increase in the metabolic rate and thermoregulation. Brown fat tissue, accumulated during late gestation, plays an important metabolic and thermal role during the first days of life. Increased mechanical load and movement impacts on the skeleton of the newborn, where the osteogenic program will also be affected by the increase in oxygen supply. The establishment of the intestinal microbiome and its crosstalk with the still immature immune system will impact as well on the program of hematopoiesis in the bone marrow which started to build at late fetal stages (Hillman et al., 2012; Liggins, 1994; Morton and Brodsky, 2016; Sanidad and Zeng, 2020). The skeleton has a pivotal position in many of these processes, given its roles in body support, movement, organ protection, hematopoiesis and hormonal and metabolic control.

Endochondral bone-derived mesenchymal progenitors are able to engage in chondrogenic, osteogenic and adipogenic programs of differentiation, in addition to their specialization into bone marrow stromal cells (BMSC) that support hematopoiesis (Bianco and Robey, 2015). Development of long bones is a dynamic and tightly orchestrated process which reflects the capacity of mesenchymal progenitors to first generate cartilage, which will serve as a mold and will provide the signals to start the program of osteogenesis and the formation of the bone marrow (BM) (Galea et al., 2021; Haraguchi et al., 2019; Long and Ornitz, 2013). During the formation of the BM, some mesenchymal progenitors will stay as BMSC, supporting hematopoiesis, but will also hold the capacity to differentiate into osteoblasts and adipocytes (Comazzetto et al., 2021; Jeffery et al., 2022; Shu et al., 2021). Genetic cell-fate tracking, prospective immunophenotype characterization and, recently, scRNAseq studies in the mouse are revealing the complex nature of the bone mesenchymal compartment, as well as the versatility of the different populations and their division of labor (Li et al., 2022a). For example, the capacity of hypertrophic chondrocytes to transdifferentiate into osteoblasts is now well accepted in the field (Hallett et al., 2021; Yang et al., 2014; Zhou et al., 2014). Moreover, bone fracture and fingertip regeneration mouse models, along with studies on limb regeneration in other vertebrates, are challenging ingrained concepts and pointing to alternative models by which different populations (e.g. fibroblastic cells) can be recruited and reprogrammed in a highly plastic fashion, which includes cellular phenotypic convergence (Jeffery et al., 2022; Johnson et al., 2020; Lin et al., 2021; Matsushita et al., 2021; Newton et al., 2019; Storer et al., 2020; Zhou et al., 2014). Most of these studies have nevertheless focused on the adult bone, and we still lack fundamental knowledge on the heterogeneity, relations and interactions between the mesenchymal and hematopoietic compartments at perinatal stages. To capture the main key features of these processes, we have generated a comprehensive scRNA-seq cellular map of all mouse endochondral bone compartments just before and after birth (E18.5 and postnatal day [PN] 1). The analysis of these datasets, in comparison with those from adult mice (Baccin et al., 2020), uncovers that the composition and molecular fingerprint of several bone populations change significantly between these stages. Of note, our resource study reveals the presence of distinct perinatal fibroblastic mesenchymal populations with molecular signatures that suggest their involvement in the formation of the bone marrow, organization of angiogenesis and peripheral innervation, as well as in the establishment of the environment for proper hematopoiesis, including putative direct interactions between specific mesenchymal and hematopoietic clusters. Notably, our study identifies some mesenchymal clusters with active immunomodulatory transcriptional programs which may be involved in setting an anti-inflammatory setup with implications for the maturation of the immune system and the self from non-self discrimination that is just starting to take shape. Overall, our findings highlight the relevance of integrative ontogenic studies that take into account the full cellular complexity of the endochondral bone and have important implications for the prospective isolation of cell populations for tissue engineering applications as well as for the age-tailoring of disease treatments.

## RESULTS

### A cell atlas of the perinatal endochondral bone

To get a comprehensive understanding of the composition and dynamics of the endochondral bone in the transition from intrauterine to extrauterine life, with particular emphasis on the mesenchymal compartment, we adopted a FACS-enrichment strategy that ensured that all cellular components of the E18.5 and PN1 endochondral bone (forelimbs excluding the autopod) could be captured by scRNA-seq in a balanced manner. This strategy is similar to that previously employed for adult bone (Baccin et al., 2020), and allows to investigate not only the relationships between mesenchymal subpopulations, but also their involvement in angiogenesis, hematopoiesis, and peripheral nervous system development. To represent both abundant and minor populations, we employed the sorting strategy depicted in Fig. 1A (see also Materials and Methods). Briefly, dead cells and multiplets were excluded by gating, and no lineage labeling was used to prevent the loss of any cell subsets. The gates were defined by the use of a mix of pan-antibodies to mesenchymal (CD140a/PDGFR-a) and endothelial cells (CD31/PECAM), together with CD9, a marker highly expressed in early hematopoietic progenitors and stromal cells (nicheview.shiny.embl.de; Baccin et al., 2020). Sorted cells were mixed in different proportions to better represent less abundant populations (Fig. 1A). While this strategy is not quantitative, it allows monitoring relative changes in cell number between equivalent clusters at both stages. After QC, 7,272 (E18.5) and 7,277 (PN1) high-quality cells were recovered for analysis, with a mean of 2,625 and 2,533 genes per cell, respectively.

**Fig. 1.**
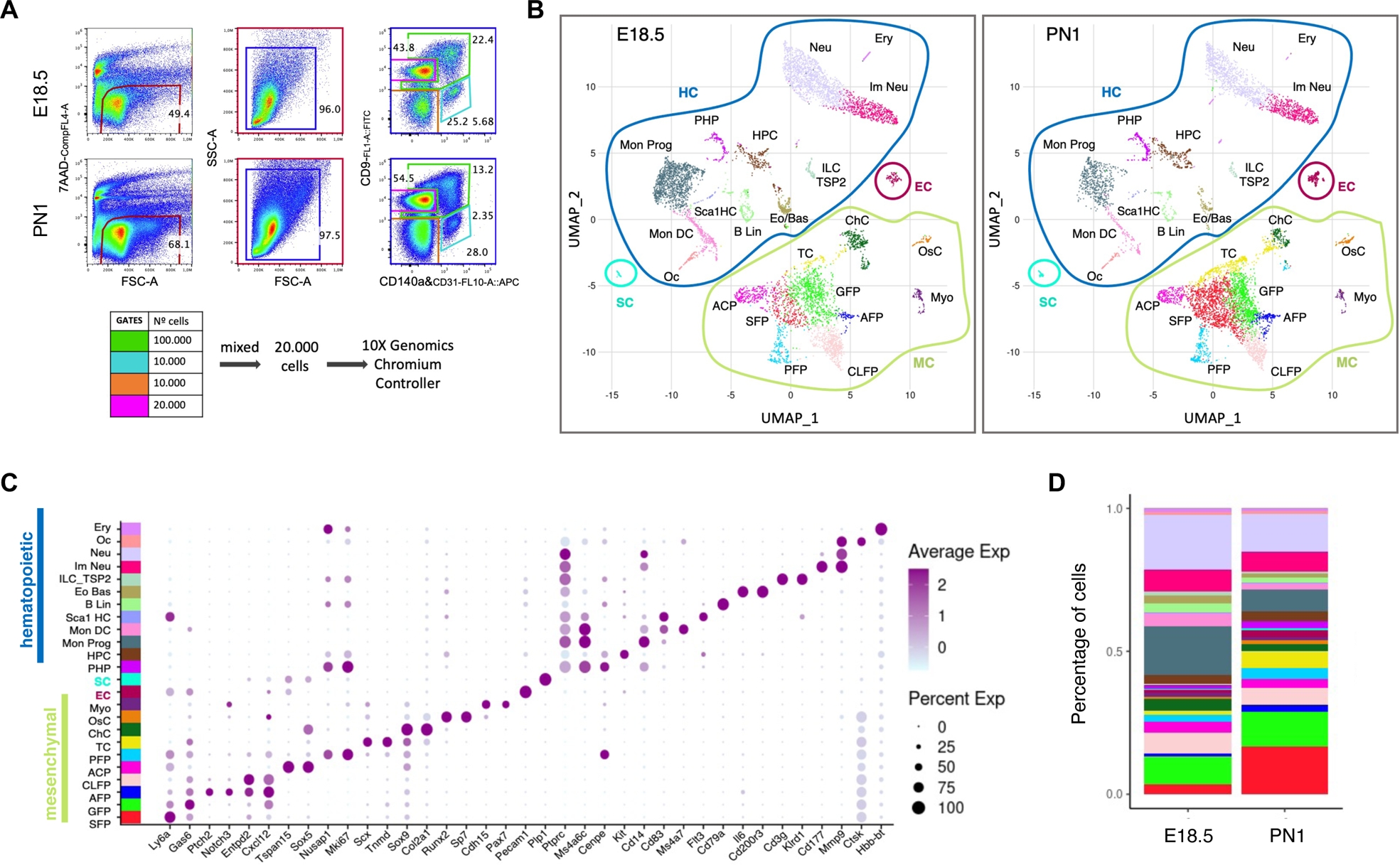
A cell atlas of the perinatal endochondral bone. (A) Gating strategy for the processing of cell suspensions for scRNA-seq, obtained after enzymatic digestion of forelimb long bones at E18.5 and PN1 (see Materials and Methods for details). The number of sorted cells that were mixed for each gate is indicated by the color codes in the FC pseudocolor plots. (B) Uniform Manifold Approximation and Projection (UMAP) of annotated clusters at E18.5 and PN1 after Harmony integration. Mesenchymal cells (MC) are encircled in green, hematopoietic cells (HC) in dark blue, endothelial cells (EC) in magenta and Schwann cells (SC) in turquoise. Neu: neutrophils; Ery: erythrocytes; Im Neu: immature neutrophils; HPC: hematopoietic progenitor cells; PHP: proliferating hematopoietic progenitors; ILC_TSP2: innate lymphoid cells and thymic seeding progenitors 2; Eo Bas: eosinophils and basophils; Sca1 HC: Sca-1^+^ hematopoietic cells; Mon DC: monocytes and dendritic cells; Oc: osteoclasts; B Lin: B-cell lineage; SC: Schwann cells; PFP: proliferating fibroblastic population; ACP: articular cartilage population; TC: tenogenic cells; SFP: Sca-1^+^ fibroblastic population; GFP: *Gas6*^+^ fibroblastic population; AFP: adipose fibroblastic population; CLFP: *Cxcl12*-low fibroblastic population; ChC: chondrogenic cells; OsC: osteogenic cells; Myo: myofibroblasts. (C) Dotplot of the representative genes across perinatal (E18.5 and PN1) clusters. Dot color and size indicate average expression levels and percentage of cells in a cluster expressing the selected gene, respectively. Gene expression is normalized and scaled to that of the entire dataset (mean=0; s.d=1), so negative values correspond to expression levels that are lower than the mean expression in the dataset. (D) Stacked-bar charts indicating the proportions of cells assigned to each cluster at E18.5 and PN1.

A combination of unsupervised and curated clustering led to the annotation of 24 cell populations, each displaying a distinct molecular signature (Fig. 1B,C, Fig. S1 and Fig. S3). These clusters encompass all hematopoietic, mesenchymal and endothelial compartments in the perinatal endochondral bone. The entire hematopoietic compartment (HC, encircled in dark blue) and each of its clusters are characterized by the expression of defining markers such as *Ptprc* (pan-HC), *Cd79a* (B-Lin), *CD200r3* (Eo-Bas), etc. Recent scRNAseq, scATAC and scProteo-genomic data have demonstrated that FACS-isolated oligopotent progenitors represent heterogeneous mixtures of progenitor populations (Buenrostro et al., 2018; Ranzoni et al., 2021; Triana et al., 2021). Not being our primary focus, we opted for broad categories for the definition of HC clusters. For instance, HPC (hematopoietic progenitor cells) encompasses granulocyte/monocyte progenitors, megakaryocyte progenitors and LMPP (Lymphoid Myeloid Primed Progenitors); likewise the B-Lin cluster includes all stages of B cell maturation. The endothelial compartment consists of a single cluster (EC; encircled in purple in Fig. 1B), defined by the expression of pan-endothelial genes as *Cdh5 or Pecam1*/CD31.

Within the mesenchymal compartment (MC; encircled in light green), clusters corresponding to fate-committed progenitors or to differentiated cells were readily identified according to their molecular signature (Fig. 1B, C, Fig. S1 and Fig. S3). The chondrogenic (ChC) cluster was defined by the expression of the *Sox9*/*Sox5*/*Sox6* trio, which drives the transcription of *Col2a1, Acan, Col9a1, Col9a2* and *Col9a3* (Akiyama and Lefebvre, 2011). The osteogenic (OsC) cluster was labelled by the expression of *Runx2*, *Osterix*/Sp7, *Bglap*/Osteocalcin, *Spp1*/Osteonectin and *Isbp*/bone sialoprotein (Komori, 2022). Finally, the myofibroblast (Myo) cluster was characterized by the expression of key myogenic genes such as *Pax7*, *Myf5, Msc, Myod1* and *Acta2/aSMA* (Alonso-Martin et al., 2016). In addition, we defined seven closely-associated clusters of fibroblastic nature. These included a cell population with tenogenic characteristics (TC) (Jelinsky et al., 2010; Murchison et al., 2007; Rees et al., 2009; Shukunami et al., 2018; Zhang et al., 2017), expressing *Scx*/Scleraxis, *Tnmd*/Tenomodulin, *Kera/*Keratocan and *Cpxm2*, and an articular cartilage progenitor (ACP) cluster expressing *Sox5*, *Gdf5, Pthlh*, *Barx1*, *Prg4* and *Wnt4* (Fig. 1C, Fig. S1 and Fig. S3) (Chijimatsu and Saito, 2019; Pacifici et al., 2000). We also identified *Tspan15* and *Ackr2* as novel ACP markers, with implications in bone growth and remodeling (Eschenbrenner et al., 2020; Lima et al., 2017; Mizuno et al., 2020). The remaining five mesenchymal clusters were even closer in the UMAP space and annotated as GFP (*Gas6^+^* Fibroblastic Population, also enriched in *Eln*-expressing cells), SFP (Sca-1*/Ly6a*^+^ Fibroblastic Population, expressing other markers such as *Ly6c1*), AFP (Adipogenic Fibroblastic Population, expressing *Ptch2* and *Notch3*), CLFP (*Cxcl12*-Low Fibroblastic Population, expressing *Ly6h* and *Lpl*) and PFP (Proliferating Fibroblastic Population) (Fig. 1C, Fig. S1 and Fig. S3). The PFP population is composed of cells in S or G2/M phases of the cell cycle and expresses mitogenic genes such as *Mki67, Nusap1, Cenpe* and *Ccna2* (Fig. 1c, Fig. S1 and Fig. S3). PFP contains the proliferating fractions of the GFP, SFP, AFP and CLFP clusters, but minimally of ACP, which fits the long-lasting quiescency of articular progenitors (Decker et al., 2017). Cell-cycle analysis also revealed small proliferating subsets within other hematopoietic and mesenchymal clusters, including ChC and Myo (Fig. S3B and 3C). The comparison of cluster ratios showed changes in both hematopoietic and mesenchymal compartments accompanying the transition to extrauterine life, including a marked increase in the representation of SFP, AFP and TC clusters at PN1 when compared to E18.5 (Fig. 1D). Since each sample per stage was obtained by pooling littermate tissue from both sexes, we deconvoluted the scRNA-seq datasets according to the expression of sex-specific genes. All clusters annotated in the pooled samples were identified in both females and males and the trends observed in the SFP, AFP and TC clusters between E18.5 and PN1 were maintained (Fig. S2). Given these results, all downstream analyses were performed using the pooled sample.

### Highlighting the differences between perinatal and adult bone mesenchymal compartments

Next, we used Harmony (Korsunsky et al., 2019; Luecken et al., 2022) to integrate our perinatal scRNA-seq datasets with those previously reported for adult (Baccin et al., 2020), generated from femurs, tibiae, hips and spines from 8-12 weeks old females (Fig. 2, Fig. S4 and Fig. S5). This analysis indicated a good broad correlation between both stages, with related clusters falling into equivalent positions within the integrated UMAP space, with the exception of PFP, which was disconnected from the main fibroblastic clusters and split into three components (asterisks in Fig. 2A). Cell cycle analysis showed that, in contrast to perinatal stages where all compartments are proliferating, only the adult hematopoietic compartment displays cells in S and G2/M phases (Fig. 2B). Focusing on the mesenchymal compartment, adult chondrocytes and myofibroblasts had perinatal counterparts. The closely associated SFP, GFP, AFP and CLFP perinatal clusters localized in similar UMAP coordinates as the adult endosteal, arteriolar and stromal fibroblasts, while perinatal ACP and TC were not clearly identified in the adult scRNAseq dataset (see Discussion).

**Fig. 2.**
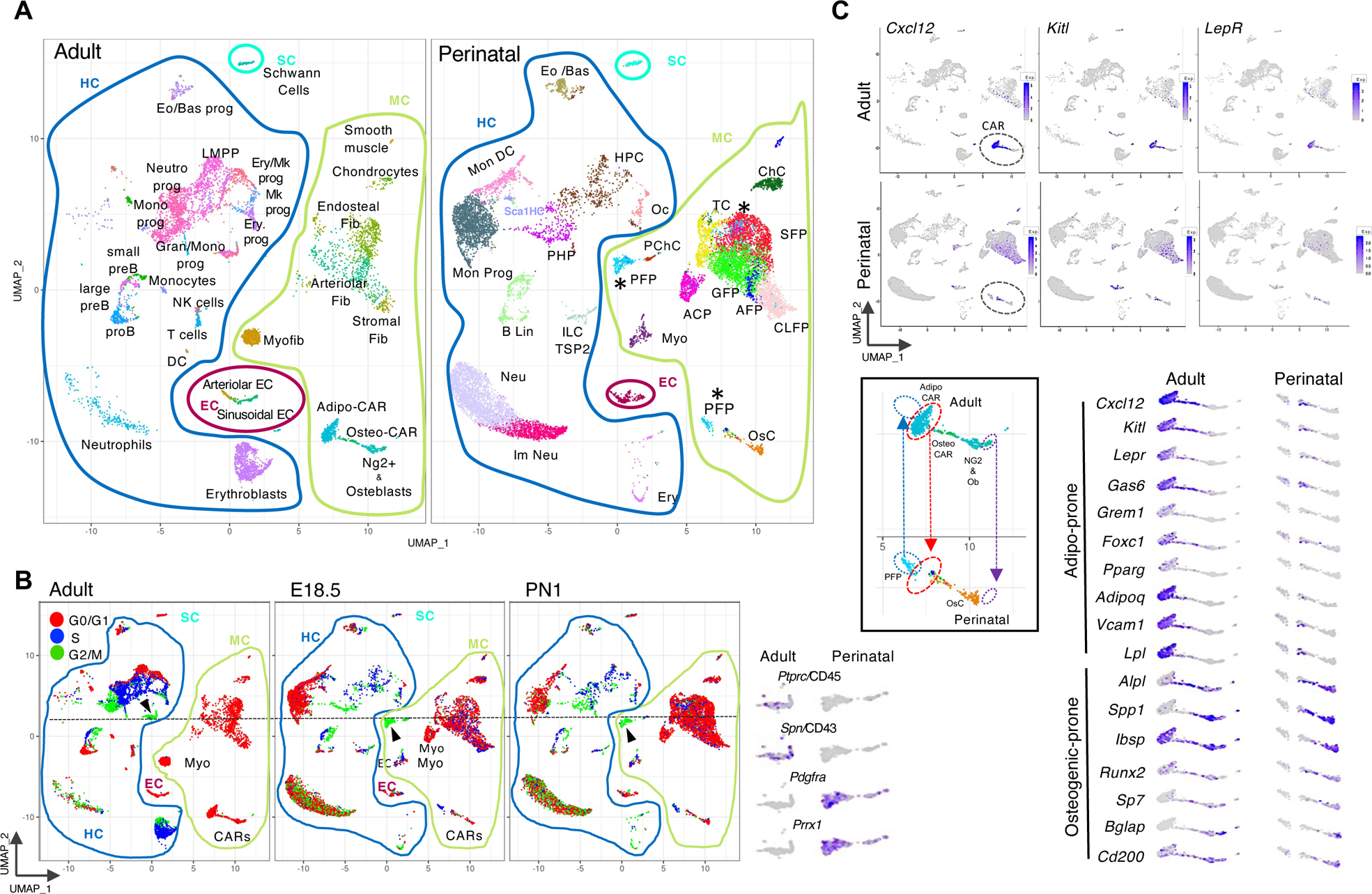
Differences between perinatal and adult bone. (A) UMAP plot of Harmony-integrated adult (Baccin et al., 2020) and perinatal scRNAseq datasets. Note that the integration with adult data changes the distribution of the clusters in comparison to Fig. 1 (e.g. PFP, indicated by asterisks). PChC: proliferating chondrogenic cells. (B) Cell cycle analysis of the Harmony-integrated adult and perinatal scRNAseq datasets. Arrowheads and dotted lines indicate G2/M clusters that in adult correspond to HC (*Ptprc+* and *Spn+*) and that in perinatal stages is part of the PFP mesenchymal cluster (positive for *Pdgfra+* and *Prrx1+*). (C) Genes expressed in adult CARs and its expression at perinatal stages (upper panels). Lower left box: enlarged view to compare the coordinates of AdipoCARs in adult and their absence in perinatal stages. Lower right panel: magnification of CAR clusters and overlay of adult Adipo- and OsteoCARs markers with the perinatal bone cellular map.

One of the most studied bone mesenchymal populations are CARs (*Cxcl12*-abundant reticular cells; Sugiyama et al., 2006). Seminal studies in the last decade have identified their pivotal role in hematopoiesis (Morrison and Scadden, 2014). Adult CARs are also characterized by the expression of *Kitl* and represent almost all LepR+ cells(Fig. 2C) (Ding and Morrison, 2013; Ding et al., 2012). scRNAseq studies have further subdivided CARs in view of their adipogenic (Cxcl12+, Alpl-) or osteogenic profiles (Cxcl12+, Alpl+) (Baccin et al., 2020; Wolock et al., 2019) and even in more subsets (Matsushita et al., 2020; Mo et al., 2022; Tikhonova et al., 2019), reflecting their adipogenic, osteogenic and BMSC differentiation potential. The comparison between equivalent UMAP plot coordinates at perinatal and adult stages (boxed area in Fig. 2C) revealed the almost complete absence of AdipoCARs in the perinatal bone and markers highly detected in adult Adipo-CARs (*Lepr, Cxcl12, Kitl, Pparg, Adipoq and Vcam1/*CD106) (Jeffery et al., 2022; Severe et al., 2019) were expressed in very few cells at perinatal stages, next to the OsC cluster (Fig. 2C). These results suggest that Adipo-CARs are just arising at perinatal stages and are in keeping with other reports using LepR antibodies, *Lepr*-CreER lines induced in the early postnatal life and recent scRNAseq analysis of the stromal compartment at postnatal day four (Kara et al., 2023; Liu et al., 2022; Mizoguchi et al., 2014; Shu et al., 2021). Of note, *Cxcl12, Kitl, Gas6* and *Lpl* were all expressed at lower levels in fibroblastic clusters, mainly in CLFP, AFP and GFP (Fig. 2C and Fig. S4), opening the possibility that some of the Adipo-CAR progenitors could be contained within these clusters. In line with this possibility, also *Pparg* and *LepR*, important players of the adipogenic program that starts to shape perinatally, are expressed in a scattered manner in the perinatal fibroblastic clusters, including AFP cells (Fig. 2C and Fig. S4). In contrast, there is a good correlation between the perinatal OsC cluster and the adult osteogenic cells, which include Osteo-CARs, Ng2+ cells and osteoblasts, with high expression of osteogenic genes (Fig. 2C and Fig. S4). As others have described for adult bone (Baccin et al., 2020; Baryawno et al., 2019; Matthews et al., 2021), we also located the perinatal prospective mSSC (murine Skeletal Stem Cells; Chan et al., 2015), characterized by the immunophenotype CD51+ CD200+; negative for CD45, CD31 and TER119, CD90, CD105, 6C3 (Fig. S5A) in the osteogenic-related OsC cluster. The integral analysis of all bone cell populations (Baccin et al., 2020 and this study) also reveals, with high resolution, the spatial and temporal (perinatal versus adult) expression patterns of reported genetic drivers and surface markers, providing an important resource for the interpretation of previous observations using cell-fate tracing mouse models and prospective isolation strategies (Fig. S5). A key advantage of including all bone endochondral populations in scRNA-seq experiments is that it provides a rich resource for the identification of markers restricted to specific populations. To illustrate this point, we identified different genes with preferential expression in the different perinatal fibroblastic clusters, which will allow the design of more precise inducible genetic tools (Fig. S6). Stressing the relevance of performing ontogenic studies, we identified genes active at perinatal stages that are not expressed in the adult tissue.

### The mesenchymal compartment in depth

Next, we generated a new Harmony representation of only the mesenchymal clusters to better illustrate their relations at E18.5 and PN1 (Fig. 3A). As previously mentioned, we observed an increase in SFP and TC populations after birth. In the case of TC (labeled by *Scx* and *Tnmd*), two different branches can be detected. One of these is related to the chondrogenic cluster, while the other branch likely represents tenogenic precursors as is preferentially labelled by additional tendon markers such as *Mkx* and *Kera* (Fig. 3B), and highly-specifically by *Ptx4*, which makes this locus a good candidate for designing genetic tools to study tendon development/regeneration.

**Fig. 3.**
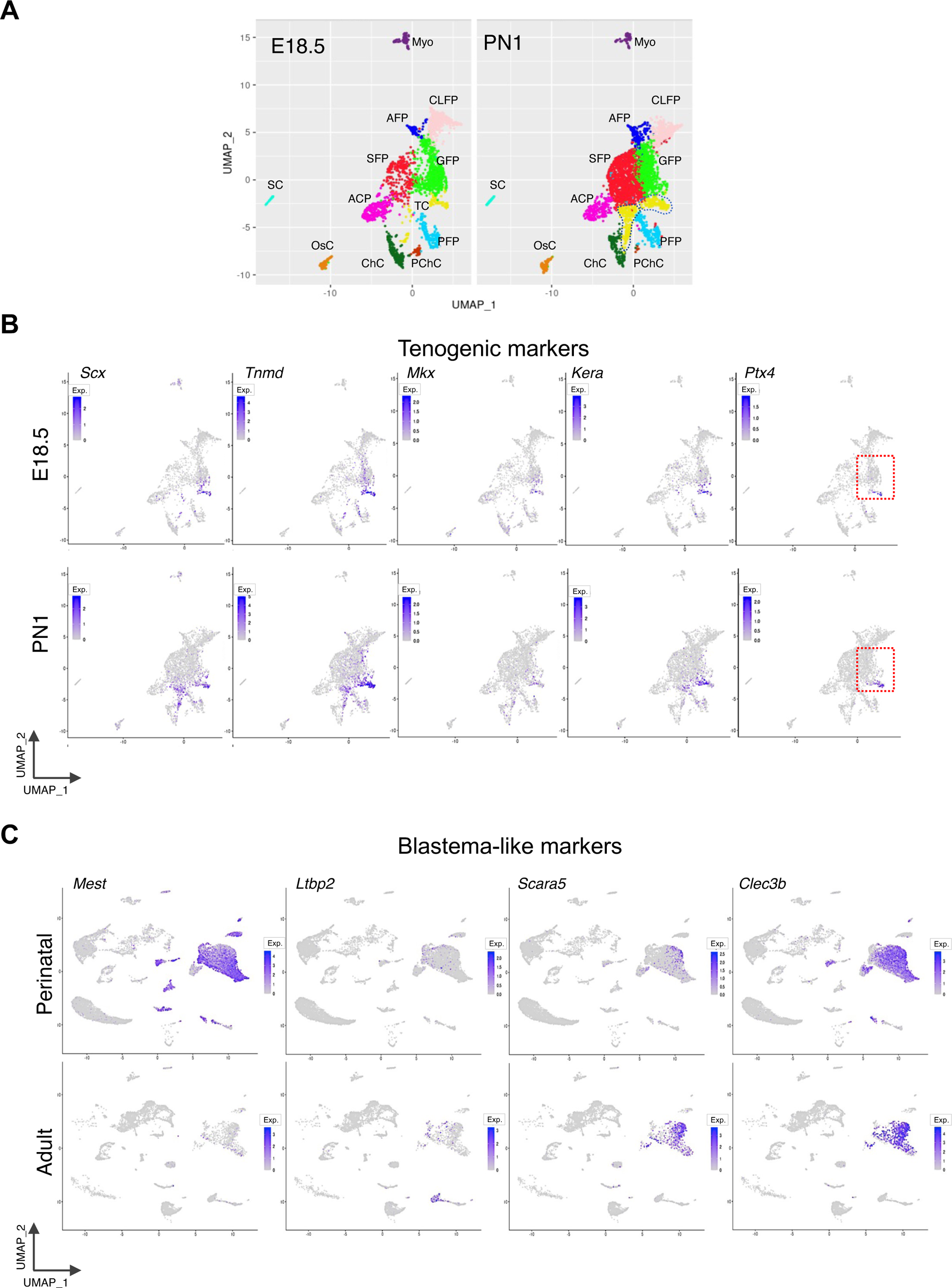
Hierarchy between mesenchymal clusters of the perinatal bone. (A) Harmony integration of mesenchymal clusters at E18.5 and PN1. The two branches of the TC cluster at PN1 are labeled by a dotted line. (B) Expression of TC-specific genes at E18.5 and PN1. Enclosed in a red dotted line is *Ptx4*, a gene exclusively expressed in the tenogenic branch. (C) Expression in perinatal and adult bone of genes identified in dermal and blastema cells in the mouse digit tip regeneration model.

To get insight into the closely connected GFP, SFP, AFP and CLFP clusters, we performed Gene Ontology (GO) analyses starting from their differentially expressed genes E18.5 and PN1 (gene number ranging between 185 and 480; Table S1 and Fig. S7). These results associated CLFP, GFP and SFP to extracellular matrix (ECM) organization and angiogenesis. AFP GO terms link it with the regulation of osteoblast and fat differentiation, as well as with brown fat cell differentiation and cold-induced thermogenesis, the latter term also shared by CLFP and GFP. Fat metabolism is essential in the first hours postpartum, in preparation for starvation and thermoregulation (Hillman et al., 2012; Li et al., 2022b). AFP GO terms also associate it to myeloid and B cell differentiation, branching morphogenesis -a key feature of vessel formation, and glial cell differentiation. SFP GO terms are related to cell migration and cell adhesion, regulation of nitric oxide synthesis, leukocyte differentiation, coagulation and wound healing. GFP displays GO terms for ossification, osteoblast differentiation, response to mechanical stimulus, fibroblast proliferation, cartilage development, hormonal regulation and glucose homeostasis (Morton and Brodsky, 2016). CLFP is also associated to bone mineralization, ossification, glial cell migration (along with SFP), regulation of muscle cell differentiation and axon guidance. This plethora of putative tasks suggests a pivotal role for perinatal fibroblastic clusters in the bone under construction and in the transition to postnatal life. In support for these broad organizing functions, we noted that the perinatal fibroblastic cluster signatures include genes previously identified in the blastema that forms after digit tip amputation in adult mice, including *Mest1*, *Ltbp2*, *Scara5*, *Clec3b* and *Cd34*, the latter three expressed in dermal-derived fibroblasts (Fig. 3C and Fig. S5) (Johnson et al., 2020; Storer et al., 2020).

### Interactions between endochondral cell subsets and implications for central tolerance

To gain insight into the interconnectivity of the different perinatal bone cell populations, we used CellPhoneDB, an algorithm based on the expression pattern of ligands, receptors and ECM components, therefore capturing both direct and indirect interactions (Efremova et al., 2020). The degree of interaction within mesenchymal clusters (MC-MC) and between MC clusters and EC (MC-EC) or hematopoietic clusters (MC-HC) is shown in Fig. 4A. In all three compartments, cluster interactions were higher at E18.5 than at PN1. MC-MC displayed the highest number of calls, being TC, OsC, SFP and GFP the most interacting clusters, and Myo the least. ECM components such as collagens and integrins were the most abundant predicted connectors, as expected for fibroblastic populations, with several stage-specific differences (Tables S2-S4). The type of collagens and integrin complexes was highly diverse and different clusters displayed specific profiles (Table S4), which fits the dynamic properties of the matrisome, which changes according to age and inflammatory conditions (Helbling et al., 2019). For example, a significant difference between E18.5 and PN1 was the lack of integrin a1b1 complex at PN1 in GFP, ACP and OsC clusters. This integrin complex is associated with remodeling and wound healing (Carver et al., 1995). Membrane-bound COL13A1, relevant for bone growth, was mainly expressed by OsC and EC clusters (Kemppainen et al., 2022; Ylonen et al., 2005). This analysis also identified the interconnection of SC with several other subsets through *Col20a1* and a1b1/a2b1 integrin complexes. *Col20a1* is SC-specific and expressed only at perinatal stages (Fig. S1, Fig. S3 and nicheview.shiny.embl.de). As the ECM participates in signaling, migration, lineage specification and compartmentalization in several tissues (Long and Huttner, 2019), these changes might dictate a code involved in the spatial and temporal organization of the skeletal program (Stone et al., 2023).

**Fig. 4.**
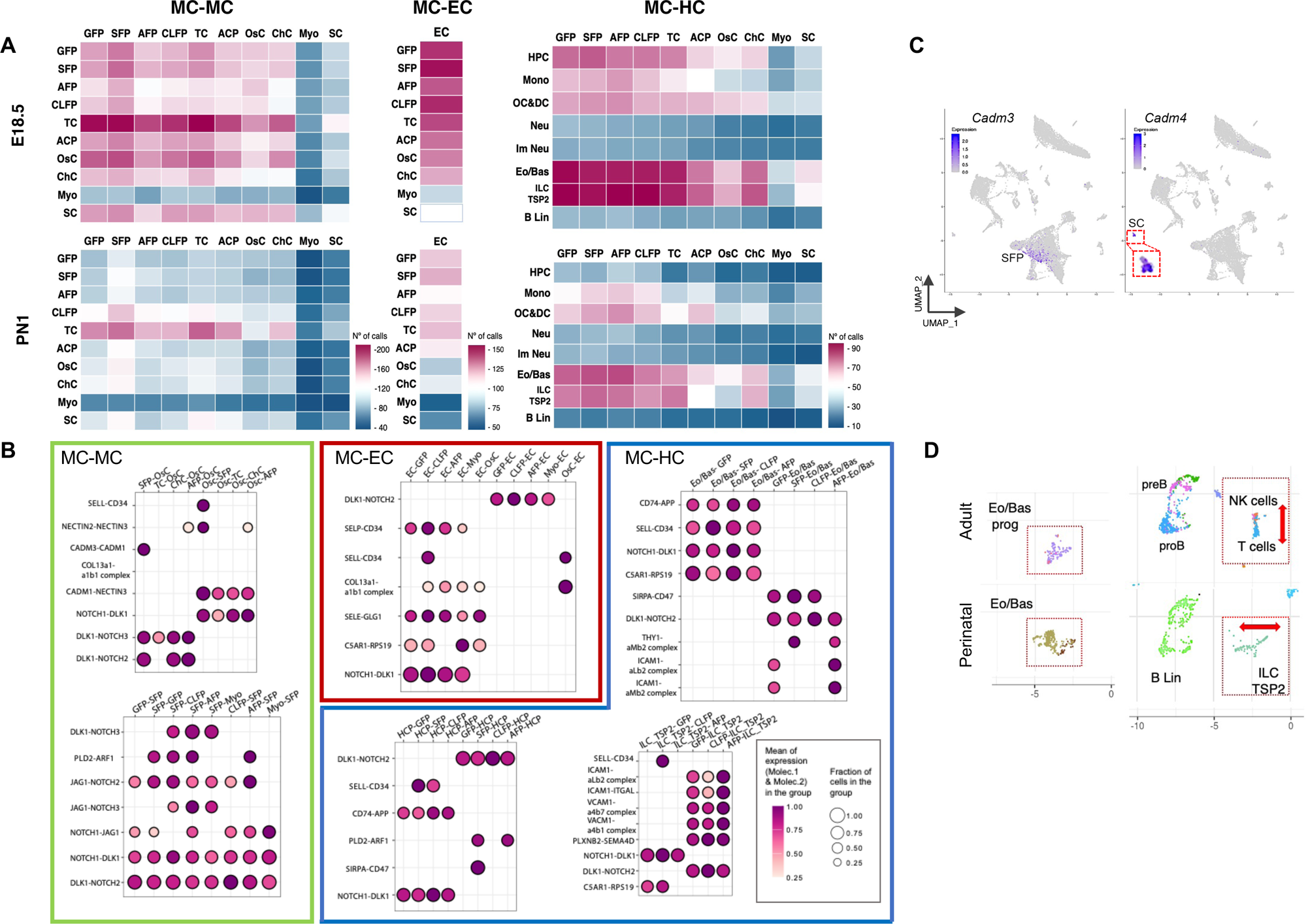
Cell to cell communication in the perinatal endochondral bone. (A) Heatmap showing the total number of connections (N° of calls; p-val≤0.05) identified by the CellPhoneDB algorithm within mesenchymal clusters (MC-MC), between mesenchymal clusters and EC (MC-EC) and mesenchymal and hematopoietic clusters (MC-HC). (B) Bubble plot representation of selected putative direct interactions (rows) inferred with CellPhoneDB (no secreted molecules; p-value ≤0.05) at E18.5 showing the mesenchymal clusters (clusters) with the higher number of associations within the mesenchymal compartment (MC-MC), with endothelial cells (MC-EC) and with hematopoietic clusters (MC-HC). Dot size represents the mean fraction of cells in the group (cluster1 and cluster 2) expressing molecule 1 and molecule 2. Color encodes the scaled mean expression levels of interacting molecules in the group. (C) UMAP projection at E18.5 showing the exclusive expression of *Cadm3* and *Cadm4* in SFP and SC, respectively, identified by CellPhoneDB as a putative direct interaction between these clusters. (D) UMAP coordinates comparing Eo/Bas, NK-Tcells versus ILC-TSP2 and B-cell clusters in perinatal and adult bone. Striking differences are observed between NK-T cells versus ILC-TSP2 (enclosed in red dotted box and indicated by red arrows).

Next, we explored direct interactions (no secreted molecules; p-value < 0.05) and focused on those clusters displaying the highest number of partners and the strongest connectors (Fig.4B; all direct connectors are provided in Tables S5 and S6). Within the MC, the SFP and OsC clusters appeared as the most dynamic, with CADM1 and COL13a1 mediating significant interactions in the latter. NOTCH and their ligands were highly expressed among fibroblastic clusters, as expected from their important roles in bone development and homeostasis (Zieba et al., 2020). DLK1-NOTCH1 and DLK1-NOTCH2 mediated very strong connectors in all mesenchymal clusters and with EC (Tables S5 and S6). DLK1 is a non-canonical ligand that inhibits NOTCH1 receptor activity (Gonzalez et al., 2015). Of note, *Dlk1* is not expressed in adult bone (nicheview.shiny.embl.de; Baccin et al., 2020), whereas at perinatal stages it is restricted and highly expressed in most of the clusters of the mesenchymal compartment. NOTCH3 mediated various putative connectors, mainly in AFP, which is relevant as *NOTCH3* mutations cause Lateral Meningocele Syndrome (OMIM #130720), characterized by several skeletal abnormalities (Zieba et al., 2020). In addition, the highly prevalent JAG1-NOTCH3 pair could play a role in vasculature remodeling, as slow-cycling LepR:Cre *Notch3*+ cells were closely associated to vasculature in the bone marrow of adult mice (Mo et al., 2022). In fact, the EC connectome (also explored at E18.5 by Liu et al., 2022), displayed a high number of direct predicted interactions with GFP, OsC, AFP, CLFP and Myo, mainly through selectins (SELL, SELP, SELE), supporting our previous GO analyses that associated AFP and CLFP with different aspects of angiogenesis (Fig. 4B and Fig. S7). Schwann cells (SC) also showed broad potential direct interactions with SFP, AFP, OsC, CLFP and with themselves (Table S5). In particular, we identified a specific link mediated by CADM3 (exclusive of SFP) and CADM4 (exclusive of SC) (Fig. 4C). Future studies will determine the relevance of this interaction, but CADM3 mutations are associated to type 2FF Charcot-Marie-Tooth disease (OMIM #619519), a peripheral neuropathy with early childhood onset and characterized by progressive weakness and muscle atrophy (Rebelo et al., 2021). In this context, both *Col20a1* and *Cadm4* are excellent candidate *loci* for the design of genetic tools to address peripheral nervous system regulation of endochondral bone development and regeneration (Johnston, 2017).

Concerning the hematopoietic compartment, several of the MC clusters were predicted to interact with HPC, and preferentially with Eo/Bas and ILC-TSP2, being GFP, SFP, AFP and CLFP the most active (Fig. 4A). Representative direct connectors for HPC, ILC-TSP2 and Eo/Bas are APP-CD74, ICAM1-ITGAL and SELL-CD34, respectively (Fig. 4B and Table S6). While adult/perinatal Eo/Bas and B lineage cells map to equivalent coordinates in the UMAP space, there are significant differences between perinatal ILC-TSP2 cells and adult NK and T cells (Fig. 4D and Fig. S8). T cells in the adult bone marrow correspond to memory T cells and regulatory T cells (Di Rosa and Gebhardt, 2016). The T cell cluster in the adult scRNA-seq dataset (nicheview.shiny.embl.de; Baccin et al., 2020) highly expresses all CD3 components of the TCR receptor (*Cd3e*, *Cd3g*, *Cd3d* and *CD247*/TCR zeta) in addition to CD8a and CD8b1, identifying them mostly as memory CD8 T cells. In contrast, perinatal ICL-TSP2 cells only transcribe *Cd3g* and *CD247*, do not express *Lef1* (Shan et al., 2021) and are negative for *Sell/*CD62L and *Ccr7* genes expressed in central memory T cells (Jung et al., 2016). In line with recent scRNAseq data on thymus seeding progenitors (Liu et al., 2021) (TSP), we named this cluster ILC-TSP2 on the basis of the expression of *Cd7, Itgb7, Irf8, CD3e (CD3g* in mouse) and the absence of *Hoxa9* and *Cd34* transcripts (Lavaert et al., 2020; Rothenberg, 2021). Of note, we identified ILC-TSP2 as the only cluster expressing *Ccl5* (Fig. S1 and Fig. S8). This is relevant because callus formation in bone fracture models depend on the expression of the Ccl5 receptors Ccr5 and Ccr3 in periosteal Mx1+Acta2/aSMA+ cells (Ortinau et al., 2019), which highlights the relevance of the crosstalk between the MC and HC (see also additional ILC-TSP2 markers in Fig. S1; Cumano et al., 2019; Zeng et al., 2019). The role of the Eo/Bas cluster at these stages is less characterized. Perinatal Eo/Bas specifically expressed high levels of anti-inflammatory interleukins *Il4* and *Il13* (Fig. S1 and Fig. S8). In keeping with an anti-inflammatory setup, we also observed the lack of expression of MHCII genes (*H2-Aa* and *H2-Eb1*) in monocyte and dendritic cells (Mon DC), and a high expression of *Il1rn* and *Il1r2* (both encoding decoy receptors of pro-inflammatory IL1) in monocytes and neutrophil clusters. In addition, monocyte, neutrophil and Eo/Bas clusters express high levels of *Osm* (Oncostatin M), which stimulates osteogenesis and inhibits adipogenesis (Li et al., 2022a), further highlighting the interdependency of the different cellular components (Fig. S8).

### Main transcriptional networks and immunomodulatory properties of the mesenchymal compartment within the perinatal endochondral bone

To shed light on the transcriptional programs operating in the mesenchymal compartment, with special emphasis on the fibroblastic clusters, we utilized SCENIC (Single-Cell rEgulatory Network Inference and Clustering) (Aibar et al., 2017; Van de Sande et al., 2020). This tool can detect the activation of gene regulatory networks controlled by a given transcription factor [called “regulons”, noted with a (+) symbol] based on the expression of its cognate targets, even if the upstream TF was not captured in the scRNA-seq data. Conversely, SCENIC labels regulons as not active if not sufficient targets are expressed, even if the controlling TF is detected. The top-ten regulons at E18.5 and PN1 and the differentially active intrauterine and extrauterine regulons are shown in Fig. S9. SCENIC accurately captured biologically relevant regulons in chondrogenic [Sox9(+), Sox5(+); ChC and ACP], osteogenic [Runx2(+), Sp7(+), Dlx5(+); OsC] and adipogenic [Pparg(+), Gata6(+), Prdm6(+); AFP and GFP] clusters (Fig. S10). Concerning the latter differentiation program, *Pparg* is a key adipogenic regulator, while *Gata6* has been recently associated to brown adipogenesis (Rao et al., 2022), in agreement with our GO analysis. Finally, GWAS studies have linked *PRDM6* to obesity and osteoporosis (Hu et al., 2018), and another family member, *Prdm16*, has been implicated in brown fat differentiation in early myoblast progenitors (Seale et al., 2008; Van Nguyen et al., 2020). The CLFP population shared several active regulons with the Myo cluster [MyoG (+), Msc(+), Myf5 (+) and MyoD (+)], suggesting that these two populations could be related. Mkx(+) was the most-significant regulon in TC (Fig. 3) (Liu et al., 2015; Milet and Duprez, 2015). Two additional key observations could be extracted from the SCENIC analysis. Firstly, several Hox regulons were active in GFP, SFP, TC and ACP clusters (Fig. S9). While *Hox* functions have been most studied in the context of axial and limb patterning (Mallo, 2018), it has been described that periosteal Hox+ fibroblastic populations contribute to bone repair in adult fracture models (Rux et al., 2016). Secondly, several of the most prominent regulons in SFP and ACP were related with inflammation and showed multiple cross-regulatory interactions (tables in Fig. 5). *Nfkb1*, one of the most prominent regulons detected by SCENIC, encodes p105 and its processed form p50. Dimers of p65 (the class 2 subunit of NF-κB transcription factor, encoded by *RelA*) and p50 trigger a pro-inflammatory response. The NF-κB pathway is negatively controlled by p50/p50 homodimers that outcompete p65/p50 dimers and also suppress pro-inflammatory gene expression by association with other proteins like HDACs and p300 (Cartwright et al., 2016). p50/50 homodimers can also promote the expression of anti-inflammatory genes by association with Bcl3 (Wessells et al., 2004), and the Bcl3(+) regulon itself is active at E18.5 and PN1 and controls *Nfkb1* expression (Fig. 5 and Fig. S9). Another active regulon involved in inflammation regulation and active at E18.5 and PN1 is Atf3(+) (Hai et al., 2010). ATF3 has been shown to inhibit NF-κB pathway by dimerization with p65 and recruiting HDAC1 (Kwon et al., 2015). *Klf3* is a transcriptional repressor that directly suppresses the expression of *RelA* (Knights et al., 2020), and its regulon is active only at E18.5. Of note, the Klf4(+) regulon is detected as active at PN1. In addition to its role in osteogenesis (Abe et al., 2022), *Klf4* participates in several cellular processes depending on the cellular type and context (Ghaleb and Yang, 2017) and it can act as either a pro- or anti-inflammatory factor (Choi et al., 2018; Kawata et al., 2022; Li et al., 2021; Liao et al., 2011). The Cebpd(+) regulon modulates different cellular processes (Balamurugan and Sterneck, 2013), and is mainly related with inflammatory response in macrophages but can also act in preventing deleterious effects of the inflammatory response (Moore et al., 2012; Rustenhoven et al., 2015). Other regulon active only at PN1 and associated with inflammatory regulation is Nr1d1(+) (Nr1d1/Rev-Erbα encodes a core protein of the circadian clock) (Curtis et al., 2014; Liu et al., 2020; Scheiermann et al., 2013). Interestingly, and still not explored in depth, different *Hox* genes interact with the NF-κB pathway and intervene in different aspects of inflammation (Mucientes et al., 2018; Pai and Sukumar, 2020).

**Fig. 5.**
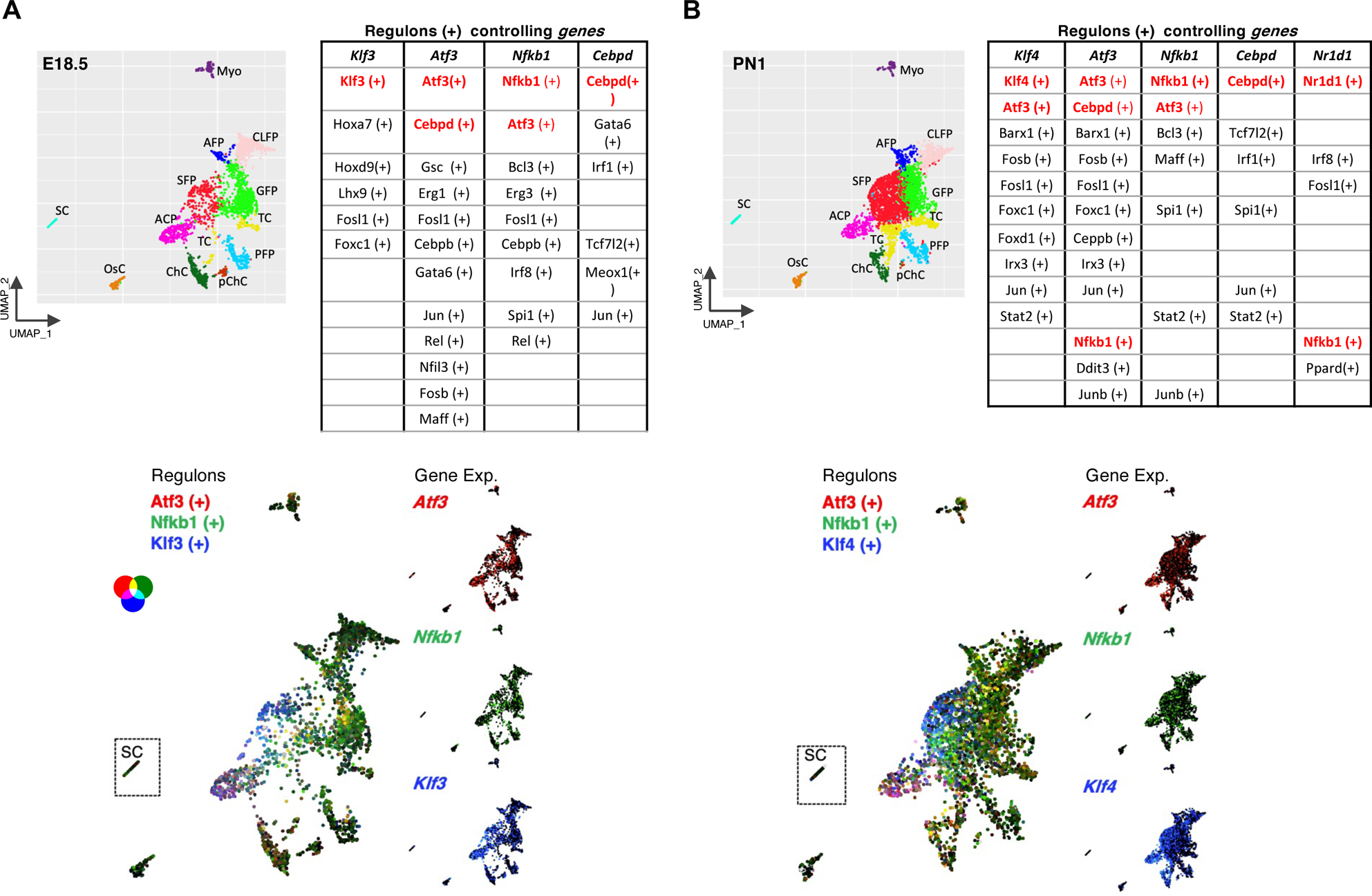
Immunomodulatory properties of the perinatal bone mesenchymal compartment. (A, B) SCENIC analysis of the MC at E18.5 (A) and PN1 (B) displaying the main immunomodulatory regulons. UMAP projections of the mesenchymal clusters at E18.5 and PN1 are shown as a guide for cluster identification. The tables show the regulons that control the expression of key transcription factors involved in immunomodulation (*Klf3*, *Klf4*, *Nfkb1*, *Cebpd* and *Nr1d1*). Colored in red are their regulons to show their autoregulation and the network of cross regulation. Lower panels depict the joint activity of the regulons Atf3(+), Nfkb1(+) and Klf3(+) at E18.5 and Atf3(+), Nfkb1(+) and Klf4(+) at PN1. The individual expression of the corresponding genes is displayed to the right of each SCENIC projection.

### Strategies for the prospective isolation of mesenchymal populations

Based on their molecular profiles, the well-characterized P⍺S population [immunophenotype: Lineage negative (CD45, TER119, CD31) and PDGFR-a+, Sca-1+] corresponds to the SFP, ACP and part of the GFP clusters. P⍺S were first described in adult bone (Morikawa et al., 2009) and these Sca-1+ (encoded by *Ly6a*) fibroblastic populations have been shown to contain multipotent progenitors able to give rise to cartilage, bone and adipose tissue (Ambrosi et al., 2017; Ambrosi et al., 2021; Morikawa et al., 2009; Nusspaumer et al., 2017). In addition, Sca1+ cells in the periosteum highly contributed to the formation of the callus in fracture models (Duchamp de Lageneste et al., 2018; Jeffery et al., 2022; Matthews et al., 2021; Mo et al., 2022; Perrin et al., 2023). P⍺S are most abundant at perinatal stages, peaking immediately after birth and being less represented in adults (Nusspaumer et al., 2017). These properties, together with the immunomodulatory profile of SFP and ACP revealed by the SCENIC analysis, make P⍺S an important target for biomedical applications. Hence, developing robust isolation strategies is key for the isolation of the human equivalent populations for use in cellular therapies, but as there is no human ortholog of the *Ly6a*/Sca-1 gene (Lee et al., 2013), additional reliable markers need to be identified. To do so, we interrogated our scRNA-seq datasets for several surface markers that were evolutionary conserved between mouse and human and validated them by FC in order to develop an alternative isolation strategy to enrich in PαS cells without resorting to the use of Sca-1 antibodies. The final strategy and best markers assayed to enrich in SFP, ACP and CLFP populations from newborn bone are shown in Fig. 6. All lineage negative *Ly6a*/Sca-1+ cells were PDGFR-⍺+ (Pa+), *Pdpn*/gp38+, *Cd55*+ and mostly *Entpd1*/CD39 negative (Fig. 6A). CD39+ Pa+ defined the CLFP population, while CD55 was highly expressed in ACP and SFP, with *Thy1*/CD90 only labeling SFP (Fig. 6A and 6B). These profiles allowed devising a strategy to purify CLFP, SFP and ACP populations in three steps (Fig. 6C). The separation between SFP (CD90+) and ACP (CD90^neg^) within CD55^hi^ cells was confirmed using Sca-1 and CD90 staining. While this strategy is suitable for perinatal stages, *Entpd1*/CD39 is hardly expressed in mesenchymal adult populations (Fig. 6A), which illustrates the importance of performing ontogenic studies and adapt the prospective immunophenotype to the specific stage under analysis.

**Fig. 6.**
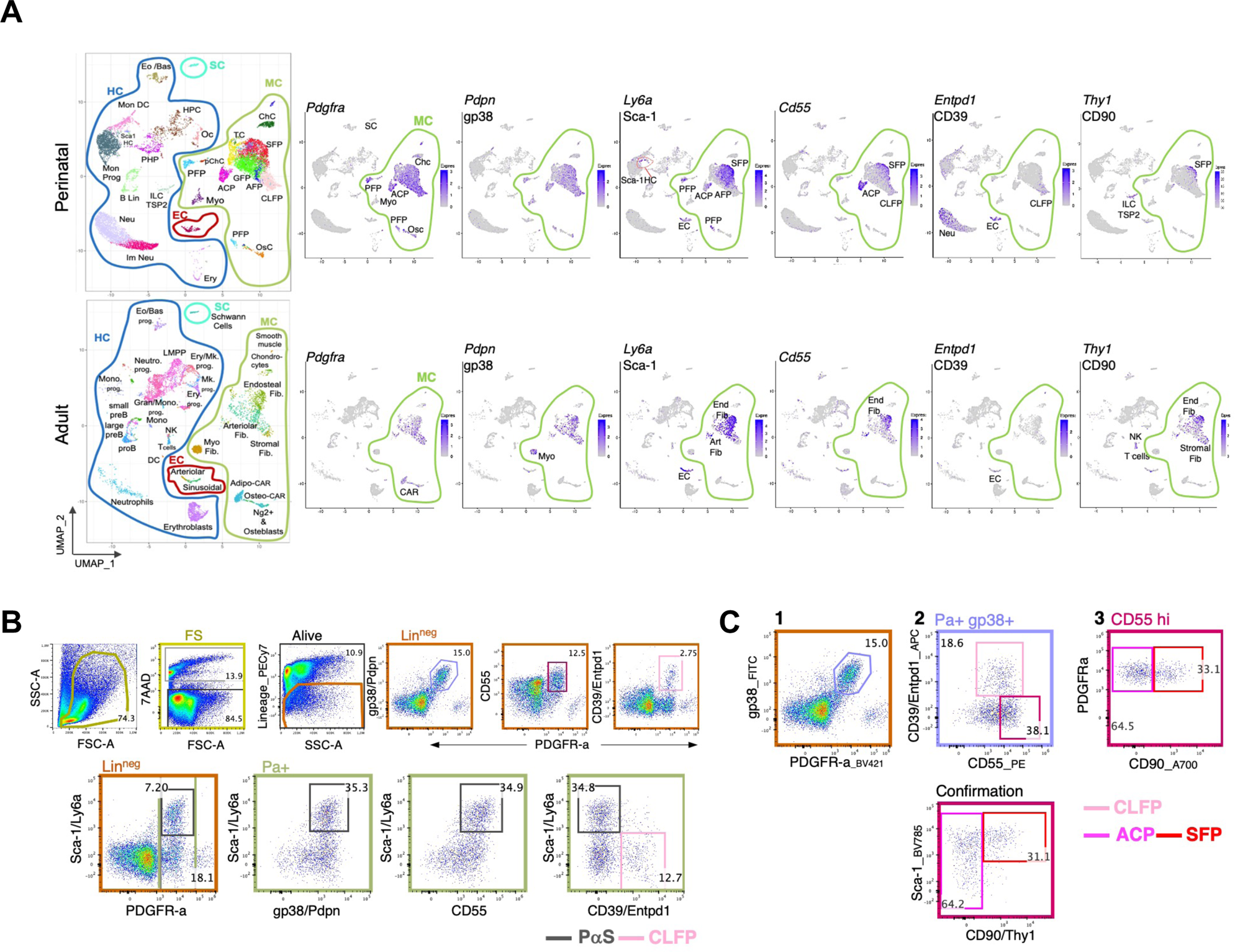
Prospective identification of ACP, SFP and CLFP clusters in newborns. (A) Expression of surface markers suitable for FC analysis at perinatal stages. PαS (PDGFRa+, Sca-1+) cells encompasses clusters SFP and ACP. In addition, *Ly6a*/Sca1 is expressed in EC and Sca1HC. *Cd55* mirrors well the expression of *Ly6a* in the mesenchymal clusters. Within mesenchymal clusters, *Entpd*/Cd39 mainly identifies CLFP, while *Thy1/*CD90 is restricted to SFP. The expression of these same genes in adult bone is shown in the bottom row. (B) FC analysis comparing the expression of Sca-1 with the other prospective markers (CD55, CD39 and CD90) in the mesenchymal compartment (Lin^neg^ PDGFR-a+/Pa+ and gp38/Pdpn+). Dead cells were excluded as 7-AAD positive. Lin= Lineage (TER119, CD45 and CD31). (C) A 3-step strategy not relying on the use of Sca-1 to enrich in SFP, ACP and CLFP populations. SFP (Lin^neg^ PDGFRa^+^ gp38^+^ CD39^neg^ CD55^+^ CD90^+^); ACP (Lin^neg^ PDGFRa^+^ gp38^+^ CD39^neg^ CD55^+^ CD90^neg^); CLFP (Lin^neg^ PDGFRa^+^ gp38^+^ CD39^pos^). Pseudocolor plots are representative of at least 3 independent experiments at PN1, each cell sample extracted from to 3-4 littermates. Similar results were obtained with E18.5 samples (data not shown).

## DISCUSSION

As previous scRNAseq studies have shown, the adult bone mesenchymal compartment is very diverse and this resource study uncovers how such complexity starts to build up in early life. Our analysis reveals different clusters of fibroblastic populations at perinatal stages, with diverse potentials as indicated by GO and gene regulatory network tools. These analyses unveiled that the most closely related fibroblastic clusters (GFP, SFP, AFP and CLFP), in addition to their potential to regulate osteogenesis, adipogenesis and chondrogenesis, may also be involved in a wide spectrum of tissue-organizing functions, including the regulation of hematopoiesis, angiogenesis, innervation, extracellular matrix organization, metabolism and hemostasis. For instance, AFP representation increased after birth and, along with the GFP population, is associated to adipogenesis, including brown fat differentiation and cold-induced thermogenesis, so important to control caloric restriction and temperature once the protection of the placenta is lost (Hillman et al., 2012). LepR+ CAR cells, highly represented in adults (Gomariz et al., 2018), were not present at perinatal stages. This is not due to the enrichment strategy used, since this population was barely detected by FC from total bone enzymatic cell suspensions using the CD106/Vcam1 Adipo-CARs marker (data not shown). Our findings are in line other studies (Kara et al., 2023; Liu et al., 2022), which show that Adipo-CARs are absent a E18.5 and just emerging at postnatal day 4. Interestingly, AFP could represent a precursor of Adipo-CARs, since it was the main and almost exclusive cluster expressing *Notch3*. *Notch3*-expressing cells targeted by LepRCre in adults showed to be multipotential slow cycling cells (Mo et al., 2022). Two other clusters that were more prominent after birth were SFP and TC. While the absence of an equivalent TC cluster in adults could be related to differences in sample preparation methods or in the origin of bones used, both TC and several of the fibroblastic clusters displayed differential molecular fingerprints previously identified as specific of the blastema during digit tip regeneration in adult mice (e.g. *Mest* expression). These results fit the previous observation that digit tip blastema cells are more similar to bone-derived mesenchymal cells at PN3 than to those of adult and largely differ from those of the limb bud (Storer et al., 2020). In amphibian limb amputation models, successful regeneration requires the dedifferentiation of connective tissue and dermal fibroblasts in order to rebuild the skeleton (Lin et al., 2021). Hence, the complex set of perinatal fibroblastic populations we have identified in this study -some of which also have dermal signatures-may act as reprogramming, specification and differentiation organizers of bone structure during these dynamic stages in which the bone marrow is under active construction to establish definitive hematopoiesis and during which osteo-chondrogenic programs are very active. The granularity of our study allowed the identification of various fibroblastic populations with distinct molecular identities and putative roles in these organizative processes, and our profiling of markers and active gene regulatory networks provides an entry point for the prospective isolation -as we show for SFP, ACP and CLFP- and *in vitro* expansion of some of these specific cell subsets for tissue engineering approaches. Besides, the identification of cluster-specific genes will allow the design of more precise inducible genetic mouse lines for cell fate tracking or cell population depletion. As the system reaches the steady state after sexual maturity, these fibroblastic populations become less abundant in the inner bone marrow, being associated to the endosteum and periosteum and obtained mainly by enzymatic digestion of the bone chips (Duchamp de Lageneste et al., 2018; Guarnerio et al., 2014). Concomitantly, BMSC (such as LepR+ CARs) become prominent (Jeffery et al., 2022; Mizoguchi et al., 2014; Mo et al., 2022; Omatsu et al., 2010; Shu et al., 2021). Only if the balance is broken, as demonstrated by bone fracture models, periosteal cells (e.g. with the P⍺S immunophenotype) and probably nearby connective tissue cells (Lin et al., 2021) reactivate the endochondral bone program for callus formation (Duchamp de Lageneste et al., 2018; Jeffery et al., 2022; Matsushita et al., 2021; Matthews et al., 2021; Perrin et al., 2023; Zhou et al., 2014).

Our integral approach to capture all endochondral bone cell populations also allowed us to predict intra- and inter-compartment interactions of all mesenchymal clusters. For instance, in this study we identified that *Cadm3*, the ortholog of the human gene mutated in Charcot-Marie-Tooth type 2FF peripheral neuropathy (Rebelo et al., 2021), is specifically expressed by the SFP population and mediates a putative direct interaction with Schwann cells via CADM4. These analyses also unveiled a potentially complex MC-HC connectome, inferring interactions of fibroblastic SFP, AFP, CLFP and GFP populations with HPC and, more unexpectedly, with the ILC-TSP2 and Eo/Bas clusters. From the immune perspective, these ILC-TSP2 interactions might play a role in the protection/maturation of thymus seeding progenitors for the establishment of central tolerance in the thymus and the generation of the first thymic regulatory T cells (Sakaguchi et al., 1982; Yang et al., 2015). This hypothesis is supported by the recent identification of two populations of mesenchymal cells essential for thymus development with prospective signatures similar to those of bone GFP and SFP clusters (Ferreirinha et al., 2022). Concerning the Eo/Bas cluster, we observed a high and exclusive expression of anti-inflammatory cytokines *Il4* and *Il13*. In line with our observations, basophil cells in newborns were shown to skew the differentiation of T cells towards Th2, which in newborns have an anti-inflammatory profile to protect intestinal microbiota and prevent tissue damage (Dhakal et al., 2015; Luo et al., 2023). Highlighting the predicted interdependency between compartments, both hematopoietic clusters also express key factors mediating bone healing (*Ccl5* (Ortinau et al., 2019) in ILC-TSP2) and osteogenesis-adipogenesis balance (*Osm* (Li et al., 2022a) in Eo/Bas). Future studies will be required to address the potential relevance of these bi-directional interactions at perinatal, puberty and adult stages.

Another important finding from our study is the identification of several immunomodulatory transcriptional programs operating in mesenchymal clusters (SFP, ACP and, to some extent, GFP) that seem to be skewed towards an anti-inflammatory response. These observations fit the previous proposal that the 2-week juvenile mouse bone provides an anti-inflammatory environment (with low expression of MHCI molecules in stromal populations) that changes to pro-inflammatory after sexual maturation (Helbling et al., 2019). In support of this, several mesenchymal clusters highly express *Dlk1*, a ligand previously shown to inhibit Notch-dependent pro-inflammatory cytokine production by macrophages (Gonzalez et al., 2015). In contrast to adult populations, perinatal macrophages neither express MHCII genes (*H2E-b1* and *H2A-a*) nor their protein products (also assessed by FC; data not shown), while neutrophils display high expression of the IL-1 decoy receptors *Il1rn* and *Il1r2.* An attractive hypothesis stemming from our results is that an anti-inflammatory environment of the perinatal bone marrow could ensure the proper balance between myeloid and lymphoid HSC lineage choice. High pro-inflammatory conditions promote HSC division and skew their differentiation towards the myeloid lineage at the expense of lymphopoiesis, potentially leading to excessive proliferation and exhaustion of HSC (Ho and Takizawa, 2022). In addition, and particularly in early life where individuals are first exposed to pathogens, primary immune organs (bone marrow and thymus) should be protected from infections and the deleterious effects of a highly pro-inflammatory milieu, to ensure the establishment of a proper self-tolerance.

Finally, our study reveals significant cellular and gene expression differences between neonatal and adult stages which implies that therapeutical interventions targeting endochondral bone populations or compartments must be adapted according to the age of the patient. While our work is limited by the lack of functional validation of the main findings and the fine spatial localization of each specific cluster we identify, it provides a valuable resource that highlights key aspects of endochondral bone development for the broader research community to further investigate in depth.

## MATERIALS AND METHODS

### Mice (Mus musculus)

Adult C57Bl/6J females and males were purchased from Envigo and housed under pathogen-free conditions according to Spanish and EU regulations. All animal experiments were designed according to the 3R principles and approved by the local Ethics Committee and authorities (license 01-08-2018-123). Animals were set in natural matings and vaginal plugs checked to time the collection of samples at E18.5 and postnatal day 1 (considering the day of birth as postnatal day 0). Individuals of both sexes were used for scRNA-seq experiments and flow cytometry studies.

### Tissue processing for flow cytometry

Forelimb long bones (humerus, radius and ulna) were dissected from fetuses at embryonic day E18.5 and pups at postnatal day 1 and carefully cleaned of surrounding tissue. Cell suspensions for flow cytometry (FC) analysis or sorting were prepared as previously reported (Houlihan et al., 2012), with the following specific modifications. For perinatal stages, bones were cut in small pieces with a scalpel and all tissue was processed (bone marrow cells were not washed out or flushed) for enzymatic digestion using collagenase D (2mg/ml in DMEM [high glucose]). Red cell lysis was not performed. The time of digestion for perinatal stages was 45-50 minutes in a water bath at 37°C with 3 rounds of gentle pipetting every 10 min to help disaggregation. Collagenase digestion was stopped in ice and by addition of ice-cold 10%FBS/HBSS+ (HBSS, 10mM HEPES, 1% Penicillin-Streptomycin). Cells in suspension were recovered to a new tube and any remaining bone pieces were transferred to a ceramic mortar and 2 to 3 steps of very gentle tapping with the pestle (20-30 times) was applied to increase cell recovery. Cells in the mortar were recovered with the addition of 10%FBS/HBSS+, filtered through a 100µm strainer and pooled with the cell suspension set aside. After centrifugation (350g for 10 min at 4°C), cells were resuspended in 2%FBS/HBSS+ for FC analysis or sorting. Before antibody staining, blocking of FcγR II/III was performed by incubation in ice with anti-CD16/CD32 antibodies for 15 min (1mg/million cells). Collection tubes for sorting were precoated for 15 minutes with 10% FBS/HBSS+ in ice, and kept at 4° C during sorting. Samples were analyzed and sorted in a SONY MA900 (lasers: 488/561 and 405/638nm). 7-AAD (7-amino actinomycin D) was the only dye used to discriminate dead cells. The lineage cocktail to analyze mesenchymal populations included CD45 (hematopoietic cells), TER119 (erythroid cells) and CD31 (endothelial cells), all conjugated to biotin and detected via secondary staining with Streptavidin-PECy7. Lowly-expressed molecules were stained with antibodies conjugated either with BV421 (not used when PB was selected), PE or APC, while highly expressed molecules were stained with antibodies conjugated either with PB (not used when BV421 was selected), FITC, BV785 or A700 fluorophores. All antibodies used are referenced in Table 1.

**Table 1.** Antibodies used in this study.

| **Table 1. Antibodies used in this study** | **Biolegend**<br>**Cat. No.** |
| --- | --- |
| FITC anti-mouse CD9 (clone MZ3) | 124807 |
| Brilliant Violet 421™ anti-mouse CD140a (clone APA5) | 135923 |
| APC anti-mouse CD140a (clone APA5) | 135907 |
| APC anti-mouse CD31 (clone 390) | 102409 |
| Brilliant Violet 785™ anti-mouse Ly-6A/E (Sca-1) (clone D7) | 108139 |
| Alexa Fluor 700 anti-mouse CD90.2 (Thy-1.2) (clone 53-2.1) | 140323 |
| FITC anti-mouse Podoplanin (clone 8.1.1) | 127415 |
| APC anti-mouse CD39 (clone Duha59) | 143809 |
| PE anti-mouse CD55 (DAF) (clone RIKO-3) | 131803 |
| Biotin anti-mouse TER-119/Erythroid Cells (clone TER-119) | 116204 |
| Biotin anti-mouse CD31 (clone 390) | 102404 |
| Biotin anti-mouse CD45 (clone 30-F11) | 103104 |

### Flow cytometry analysis

Data was acquired with a SONY MA900 using the equipment’s software and further analyzed using FlowJo v10.8.0. Representative plots of at least three independent experiments are shown.

### Generation of single cell RNA-seq datasets

Five embryos E18.5 and four newborns PN1, all littermates, were processed for cell suspension preparation as described above. No sex discrimination was done since at perinatal stages gender associated differences are not highly relevant. After eliminating dead cells and multiplets by electronical gating, the different fractions of sorted cells (100µm nozzle) were mixed in the proportions detailed in Fig. 1A. Cells were sorted in excess (∼5X more than calculated) to account for cell loss. Absence of cell aggregates was confirmed by microscopic visualization, and cell number and viability were assessed in a TC-20 cell counter (Bio-Rad) with trypan blue staining. For both E18.5 and PN1 samples, viability was > 90%. Cells were resuspended in HBSS+ 2%FBS at 800 cells/µl and 20.000 cells were loaded in a Chromium Controller G chip (10x Genomics) and processed with the Chromium Next GEM Single Cell 3’ Kit v3.1 for the generation of the scRNAseq libraries in parallel, according to the manufacturer’s protocol. Libraries were sequenced on a DNBSEQ-T7 system (PE100/100/10/10; sequencing performed by BGI). Depth of sequencing was 43,005 and 43,398 reads/cell at E18.5 and PN1, respectively (60% saturation). It should be noted that this kit captures the 3’UTR of transcripts, and therefore cannot detect alternative mRNA isoforms.

### Analysis of single cell RNA-seq datasets

#### Pre-processing of scRNAseq data

Reads were aligned to the mouse genome (mm10) with the Cell Ranger v6.0 software. Cells with fewer than 400 or more than 6,000 detected genes, or more than 10% of mitochondrial reads were excluded. For normalization, log-transformation with a scale factor of 10,000 was used.

#### Dimensional reduction and clustering

2,000 top high variable genes (HVG) were identified with the “vst” method of the *FindVariableFeatures* function of the Seurat package (version 4.0.3; Hao et al., 2021). The data were scaled and PCA (30 PCs) performed. The cells were clustered using the Louvain Method on a nearest neighbor graph using the *FindClusters* and *FindNeighbors* in Seurat. A UMAP on the PCA reduced data was performed to visualize the clusters.

#### Elimination of erythroid cells and doublets

Based on gene markers, erythroid cells were removed from downstream analyses. For doublet removal, the software DoubletFinder (McGinnis et al., 2019) was used. After filtering low-quality cells, putative doublets and erythroid cells, our dataset contained 7,272 cells with 10,656 mean number of reads and 2,625 mean number of genes for E18.5 and 7,277 cells with 9,827 mean number of reads and 2,533 mean number of genes in PN1. As both datasets were generated by pooling littermates of both sexes, we used sex-specific transcripts (*Xist, Esr1* and *Esr2* [female]; *Ddx3y, Eif2s3y, Kdm5d* and *Uty* [male]; Shay et al., 2020) to identify if any given cell was derived from a female or a male individual. More specifically, we assigned a “female” or “male” sex to a cell if it had at least one transcript read from a female or male specific transcript, respectively. If cells had at least one transcript read from both male and female specific transcripts the cell was tagged as “undetermined”. Cells without any sex-specific transcript reads were tagged as “NA”. For the E18.5 sample we identified 21% female cells, 42% male cells, 3.7% undetermined and 33.3% NA cells. For the PN1 sample we identified 42.3% female cells, 28.1% male cells, 4.4% undetermined and 25.2% NA cells.

#### Sample integration

In order to jointly analyze both E18.5 and PN1 samples, data were integrated with the Harmony software (Korsunsky et al., 2019) and projected into a shared 2D UMAP embedding. Briefly, the Harmony algorithm can integrate samples accounting for multiple experimental and biological factors using an iterative approach of four steps. 1) PCA for a low-dimensional embedding of cells from the different samples and soft clustering to assign cells to potentially multiple clusters favoring clusters with cells from multiple samples. 2) Global and sample-specific centroids are calculated for each cluster. 3) Centroids are used to compute linear correction factors. 4) Finally, it corrects each cell with a cell-specific factor that is a linear combination of sample correction factors weighted by the soft cluster assignment of the cell. For the comparison of perinatal with adult, we used Harmony to integrate both perinatal samples with the adult dataset (Baccin et al., 2020). The adult data is available to download as a Seurat object from https://nicheview.shiny.embl.de.

#### Cell type annotation

Marker genes described in the extensive literature were used for cluster identification. Although most of the cell types were annotated using a given clustering resolution, subsequent refinement was done through sub-clustering (*FindSubCluster* function of Seurat) and re-annotation.

#### Inference of cell-cell interactions

CellPhoneDB v.2 (Efremova et al., 2020) was used to infer cell-cell connections. To prepare the data for the CellPhoneDB analysis, mouse gene IDs were converted to their human orthologs and count data exported as h5ad format. CellPhoneDB statistical analysis was used to evaluate for significant interactions between predefined cell type pairs. For the quantification of the total number of interactions, we selected a p-value ≤0.05. For direct interactions (Table S5 and S6), we considered only connectors with no secreted molecules and p-value ≤0.05 and Log2 mean > −1.

#### Gene Regulatory Network analysis

Mesenchymal clusters were extracted and reprocessed as before (i.e. HVG, PCA, clustering, UMAP) and inference of gene regulatory networks and regulon analysis was performed using the pySCENIC software v.0.11.2 (Van de Sande et al., 2020) using default settings.

### Gene ontology analysis

Differentially expressed genes (DEG) for each of the mesenchymal GFP, SFP, AFP and CLFP clusters at E18.5 and PN1 were obtained using the FindAllMarkers() function in Seurat, using the default parameters (by default Seurat uses the Wilcoxon Rank Sum test for statistical testing). The test was run separately for each timepoint and the resulting DEG were filtered to include only genes with an adjusted p-value <0.005. The resulting number of genes used for GO analysis at E18.5/PN1 (Table S1) was 218/280 (AFP), 455/480 (CLFP), 185/234 (GFP) and 436/305 (SFP). GO terms for the “Biological Process” category were retrieved from http://geneontology.org, filtered by a ratio Fold Enriched/Expected ≥ 2 and manually curated. Plots were generated using the ggplot2 tool for R, representing the −log10 (p values < 0.05), and the overlap was calculated using the number of genes identified in the GO term for each cluster.

## ACKNOWLEDGMENTS

We thank the rest of the members of the groups for scientific discussions and technical help. We are also grateful to A. Franco, C. Mateos, A. López, P. López and L. Pérez for excellent mouse husbrandy as well as the rest of the CABD Core services, in particular C. Díaz (Flow Cytometry Facility).

## AUTHOR CONTRIBUTIONS

Conceptualization, G.N and J.L-R; Software, I.S.M and I.C; Formal Analysis, G.N, I.S.M, A.D.R, and J.J.T.; Investigation, G.N, A.D.R., I.S-A, A.A.C, A.F-M, A. M, A.M.B. and J.L-R; Writing – Original Draft, G.N, and J.L-R.; Writing – Review & Editing, all authors; Supervision, H.H, J.J.T, J.L-R., and G.N.; Funding Acquisition, H.H, J.J.T, J.L-R., and G.N.

## DATA AND CODE AVAILABILITY

- Single-cell RNA-seq data have been deposited in the GEO database (accession number GSE232202) and are publicly available as of the date of publication. Flow cytometry data reported in this paper will be shared by the corresponding authors upon request.
- All scripts necessary to replicate our analysis are available in the following Github repository (https://github.com/irepansalvador/stromal_cells.git) and is publicly available as of the date of publication.

