## Supplementary Figures and Supplementary Table Legends for "The cellular landscape of the endochondral bone during the transition to extrauterine life"

### FIGURE S1

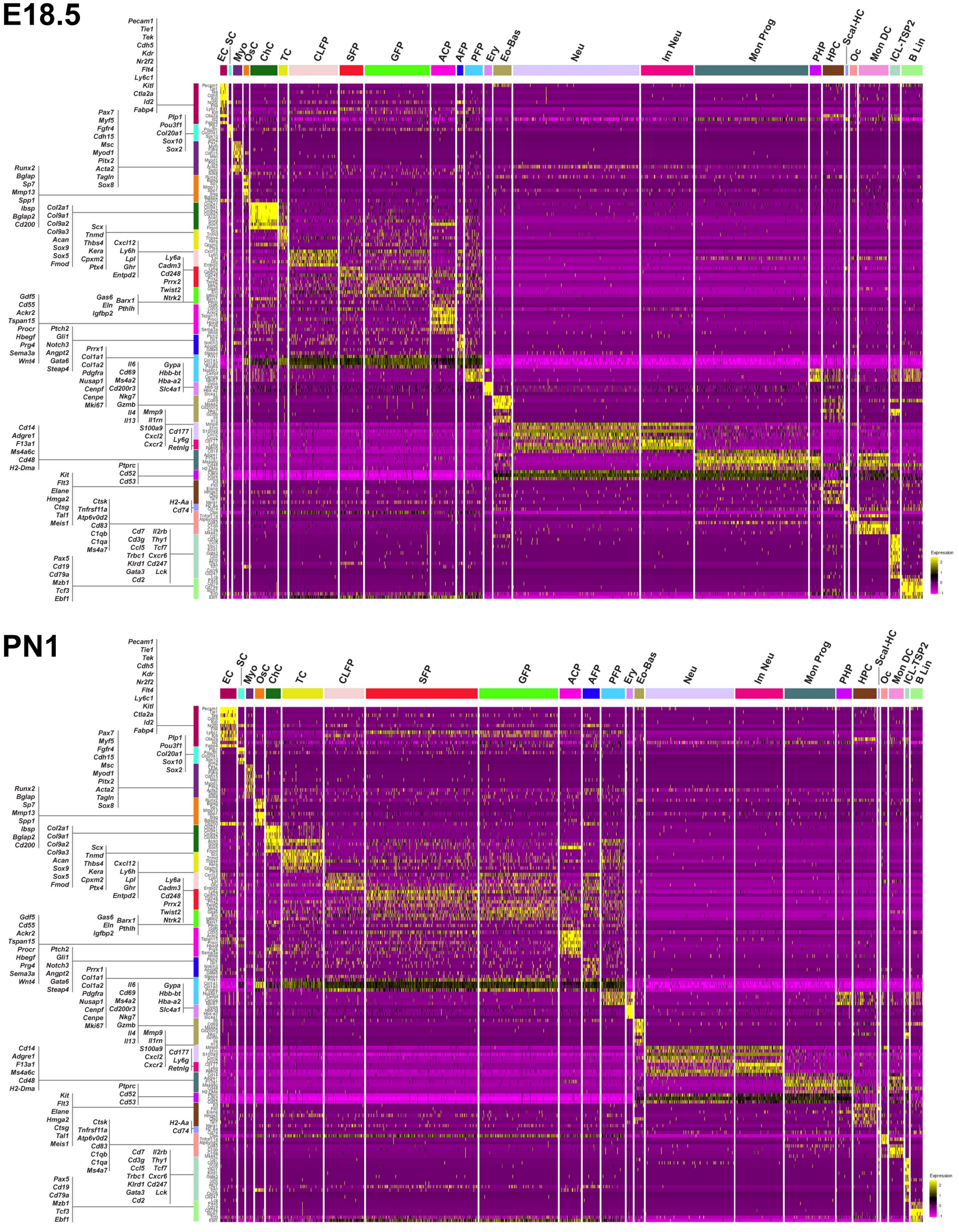


**Fig. S1. Heat map of representative cluster-enriched genes.** E18.5 (upper panel) and PN1 (lower panel). Gene expression is normalized and scaled to that of the entire dataset (mean=0; s.d=1). Negative values correspond to expression levels that are lower than the mean expression in the entire dataset.

### FIGURE S2

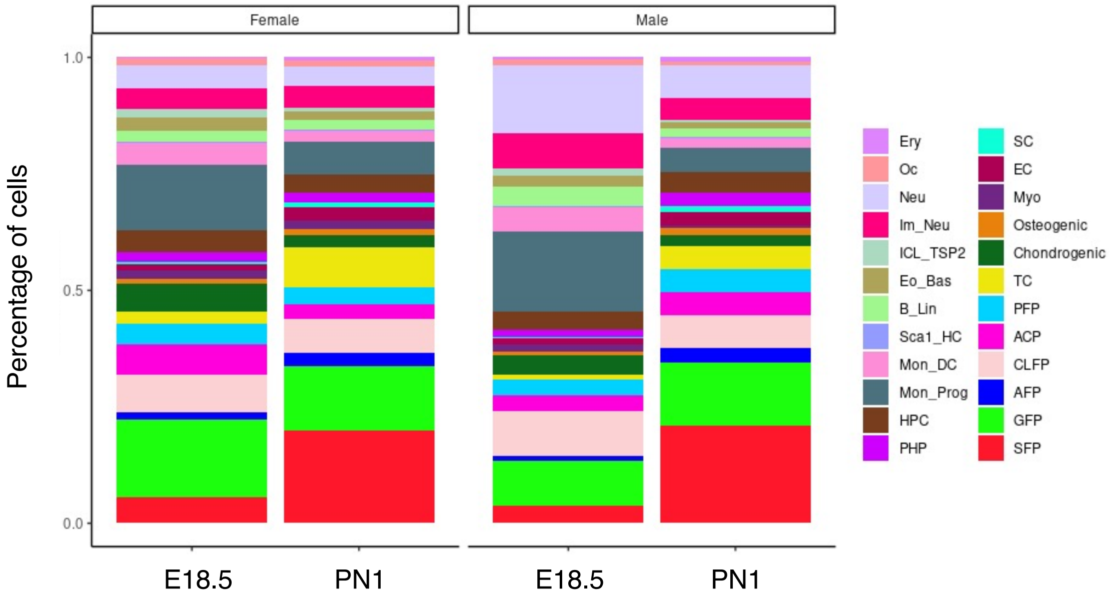


**Fig. S2. Deconvolution of perinatal bone scRNA-seq datasets according to sex.** Stacked-bar charts indicating the proportions of female and male cells assigned to each cluster at E18.5 and PN1. Note that all clusters defined using pooled samples can be also identified in both male and female replicates. Neu: neutrophils; Ery: erythrocytes; Im Neu: immature neutrophils; HPC: hematopoietic progenitor cells; PHP: proliferating hematopoietic progenitors; ILC_TSP2: innate lymphoid cells and thymic seeding progenitors 2; Eo Bas: eosinophils and basophils; Sca1 HC: Sca-1$^{+}$ hematopoietic cells; Mon DC: monocytes and dendritic cells; Oc: osteoclasts; B Lin: B-cell lineage; SC: Schwann cells; PFP: proliferating fibroblastic population; ACP: articular cartilage population; TC: tenogenic cells; SFP: Sca-1$^{+}$ fibroblastic population; GFP: *Gas6*$^{+}$ fibroblastic population; AFP: adipose fibroblastic population; CLFP: *Cxcl12*-low fibroblastic population; ChC: chondrogenic cells; OsC: osteogenic cells; Myo: myofibroblasts.

### FIGURE S3

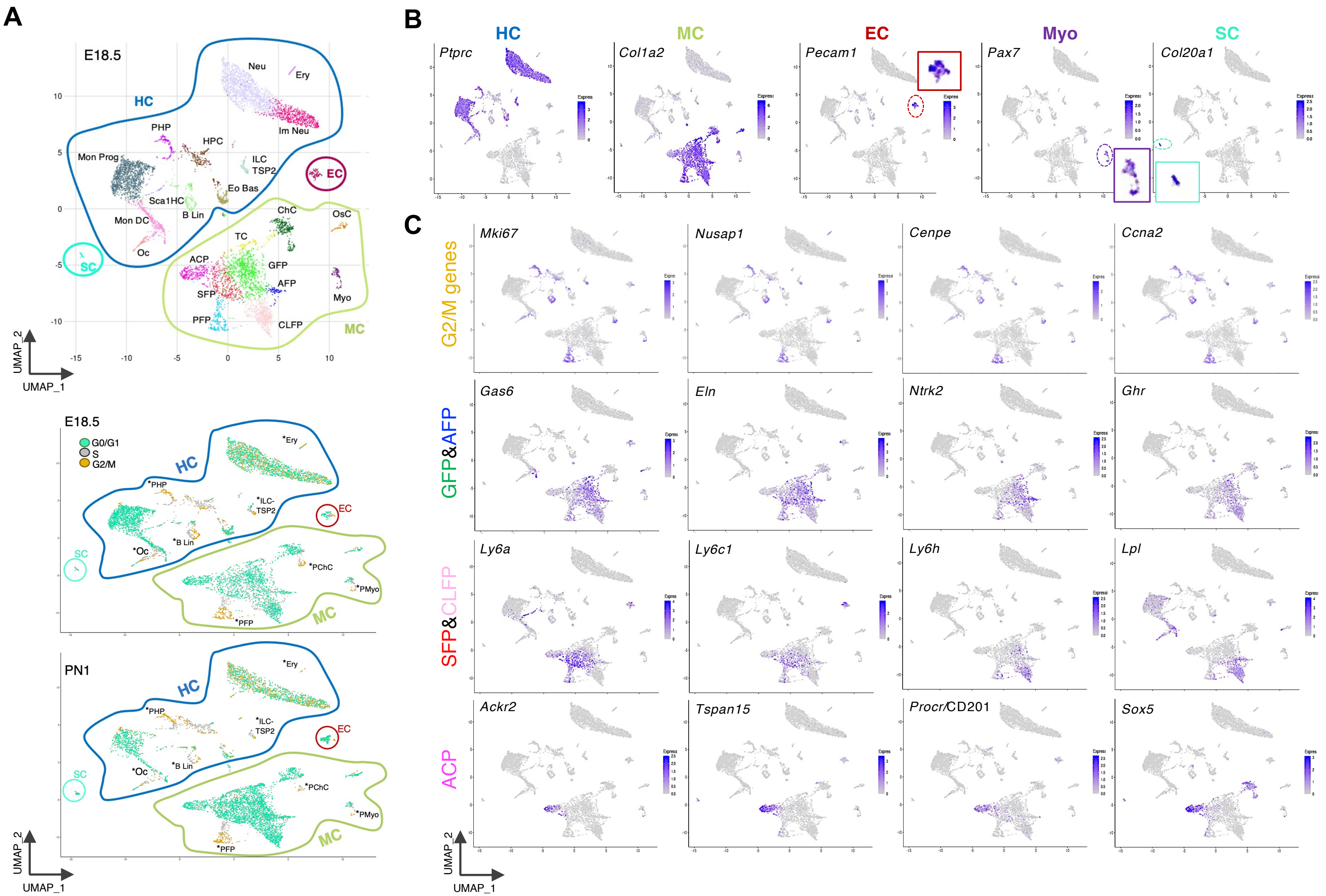
A
E18.5
HC
Neu
Ery
Im Neu
PHP
HPC
Mon Prog
ILC
TSP2
EC
Eo Bas
Sca1HC
B Lin
Mon DC
Oc
SC
ChC
OsC
TC
ACP
GFP
AFP
SFP
Myo
PFP
CLFP
MC
UMAP_2
UMAP_1
E18.5
G0/G1
S
G2/M
*Ery
*PHP
*ILC-
TSP2
*B Lin
*Oc
*PChC
*PMyo
*PFP
PN1
B
HC
MC
EC
Myo
SC
Ptprc
Col1a2
Pecam1
Pax7
Col20a1
C
G2/M genes
Mki67
Nusap1
Cenpe
Ccna2
GFP&AFP
Gas6
Eln
Ntrk2
Ghr
SFP&CLFP
Ly6a
Ly6c1
Ly6h
Lpl
ACP
Ackr2
Tspan15
Procr/CD201
Sox5

**Fig. S3. Cell cycle analysis of the perinatal endochondral bone populations.**

(A) Color coding of E18.5 and PN1 cells in the UMAP space according to cell cycle stage (bottom panel; cluster annotation of the E18.5 dataset is provided in the upper panel for reference). HC: hematopoietic compartment; EC: endothelial cells; MC: mesenchymal compartment; SC: Schwann cells. Asterisks label populations which are in S and G2/M. PChC and PMyo are proliferating chondrogenic cells and myofibroblasts, respectively. (B) Expression of compartment-defining genes and genes specific of the Myo and SC clusters. (C) Genes specific of G2/M phase (top row). The additional panels show that PFP contains proliferating cells of the fibroblastic clusters GFP, AFP, SFP and CLFP, but very few of ACP (see ACP-specific genes *Ackr2, Tspan15* and *Procr*).

**FIGURE S4**

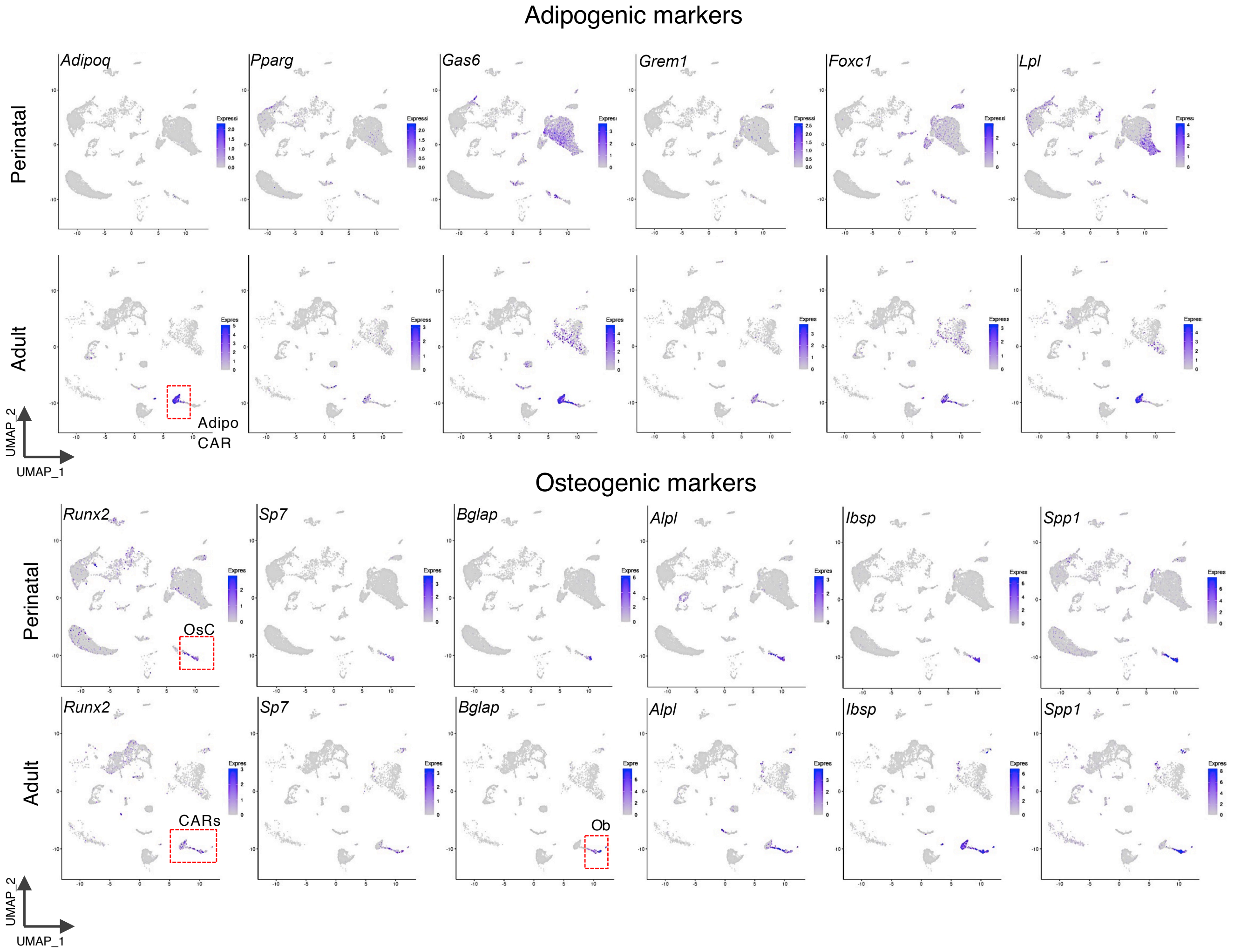


**Fig. S4. Expression profiles of adipogenic and osteogenic markers in the adult and perinatal endochondral bone.** Ob= Osteoblasts.

**FIGURE S5**

**A**

mSSC Lin Neg CD51+CD200+ (CD90, CD105, 6C3)Neg

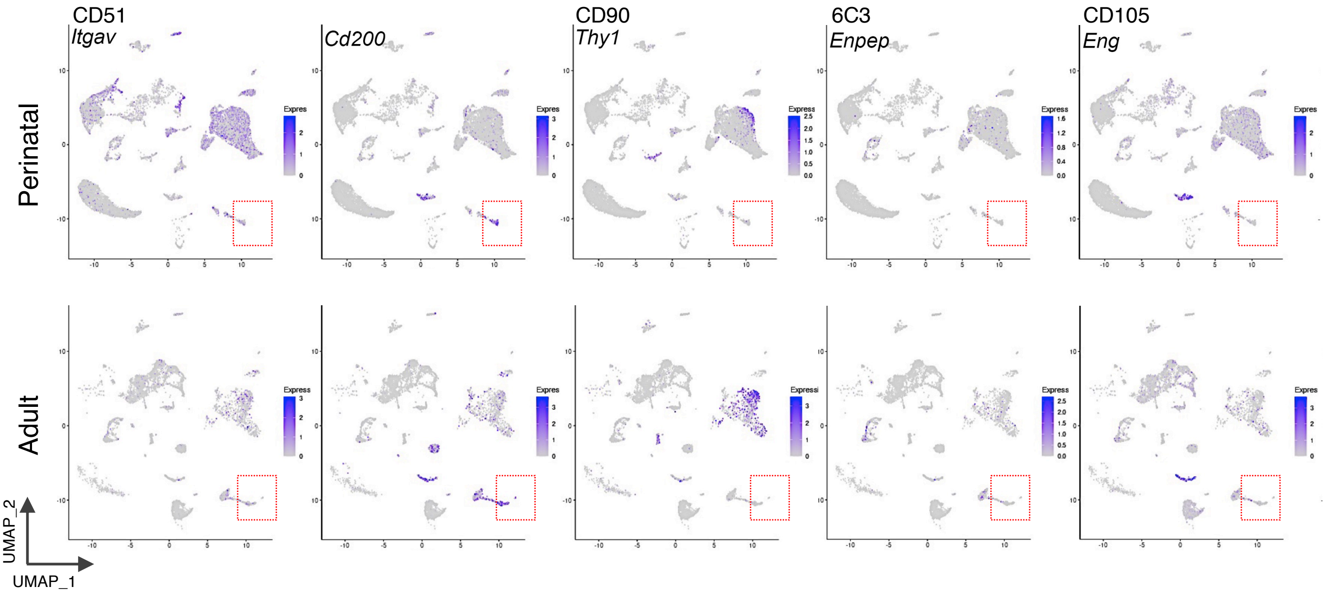


**B**

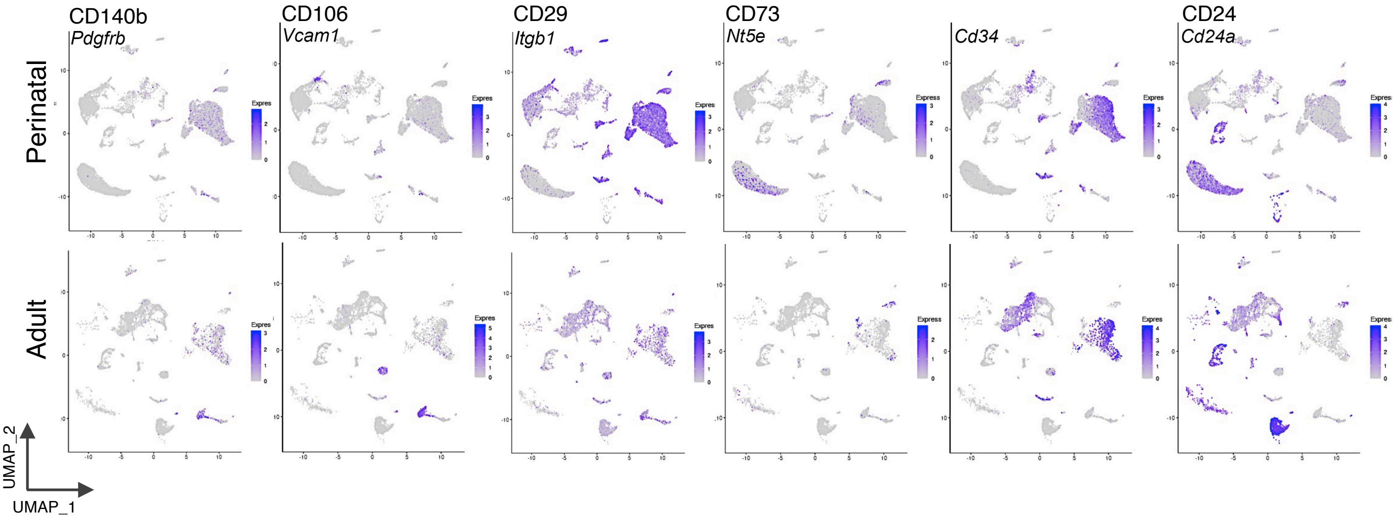


**C**

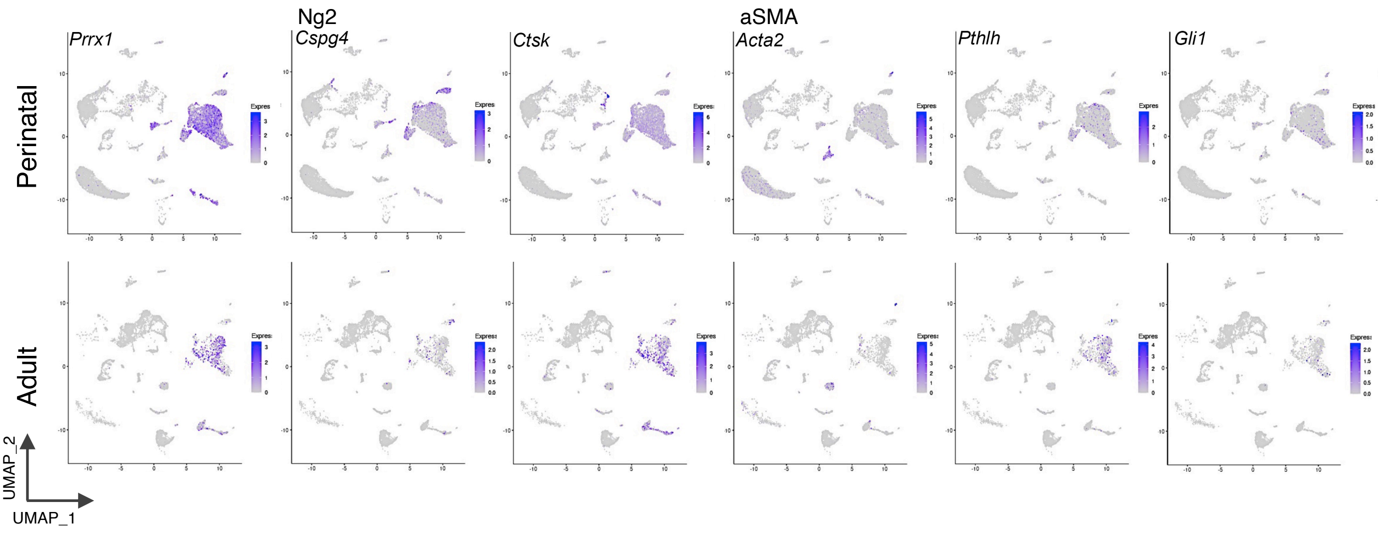

**Fig. S5. Perinatal and adult expression profile of genes used in the field for endochondral bone mesenchymal characterization.** (A, B) Expression of genes encoding surface makers used to prospectively isolate mesenchymal progenitors. Red squares indicate the cells that fulfill the mSSC immunophenotype. The advantage of prospective characterization by FC is that unwanted cells can be excluded by using the lineage strategy which is not possible using genetic approaches. (C) Expression of genes previously used in the generation of genetic tools for fate-tracking studies of mesenchymal progenitors.

### FIGURE S6

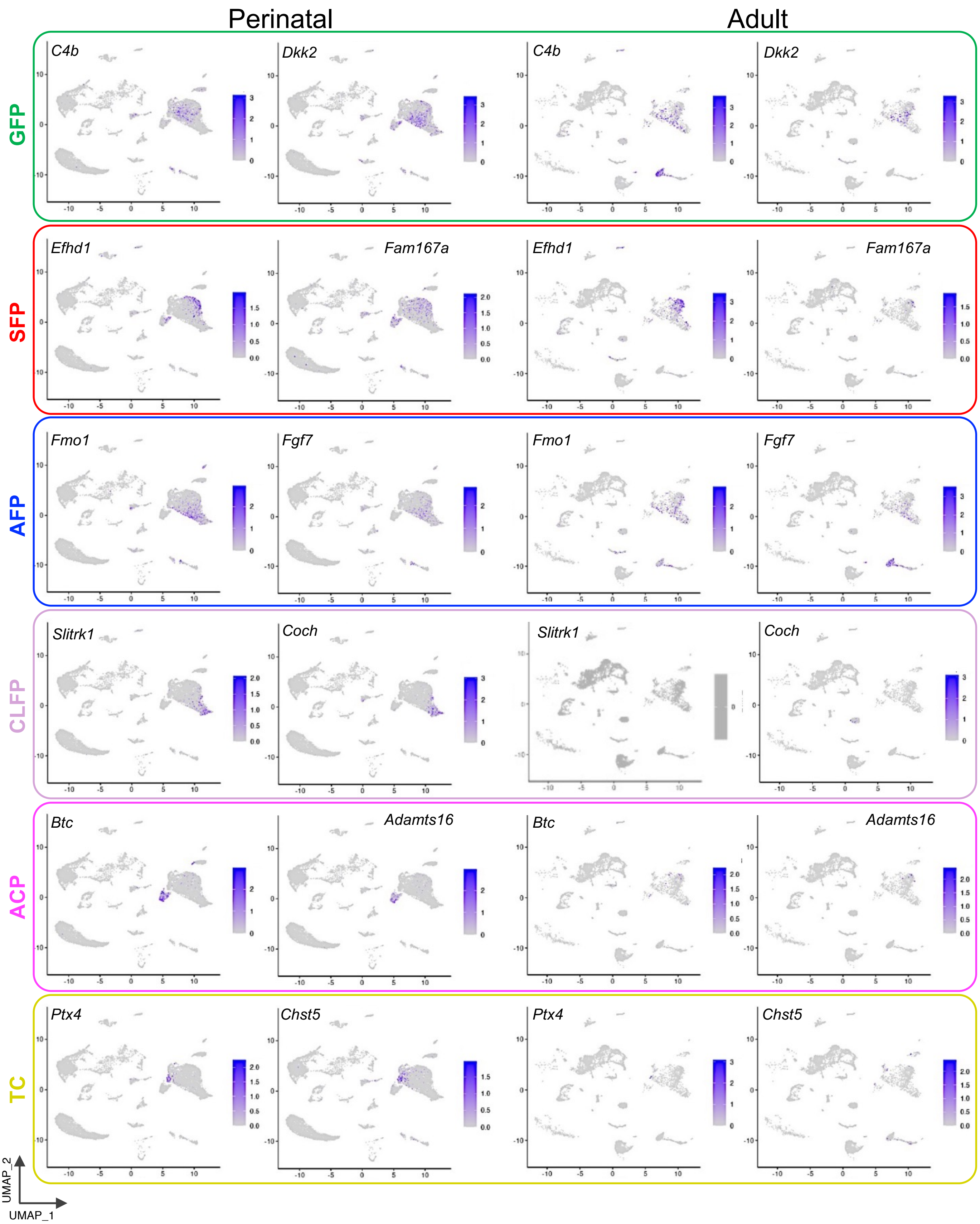


**Fig. S6. Expression of cluster-enriched markers for the generation of genetic tools.** Note the divergent expression of several genes between perinatal and adult stages (e.g. *Fam167a* in SFP cluster, *Slitrk1* and *Coch* in CLFP cluster, etc.) and that TC and ACP clusters are not represented in the adult dataset (see Discussion).

### FIGURE S7

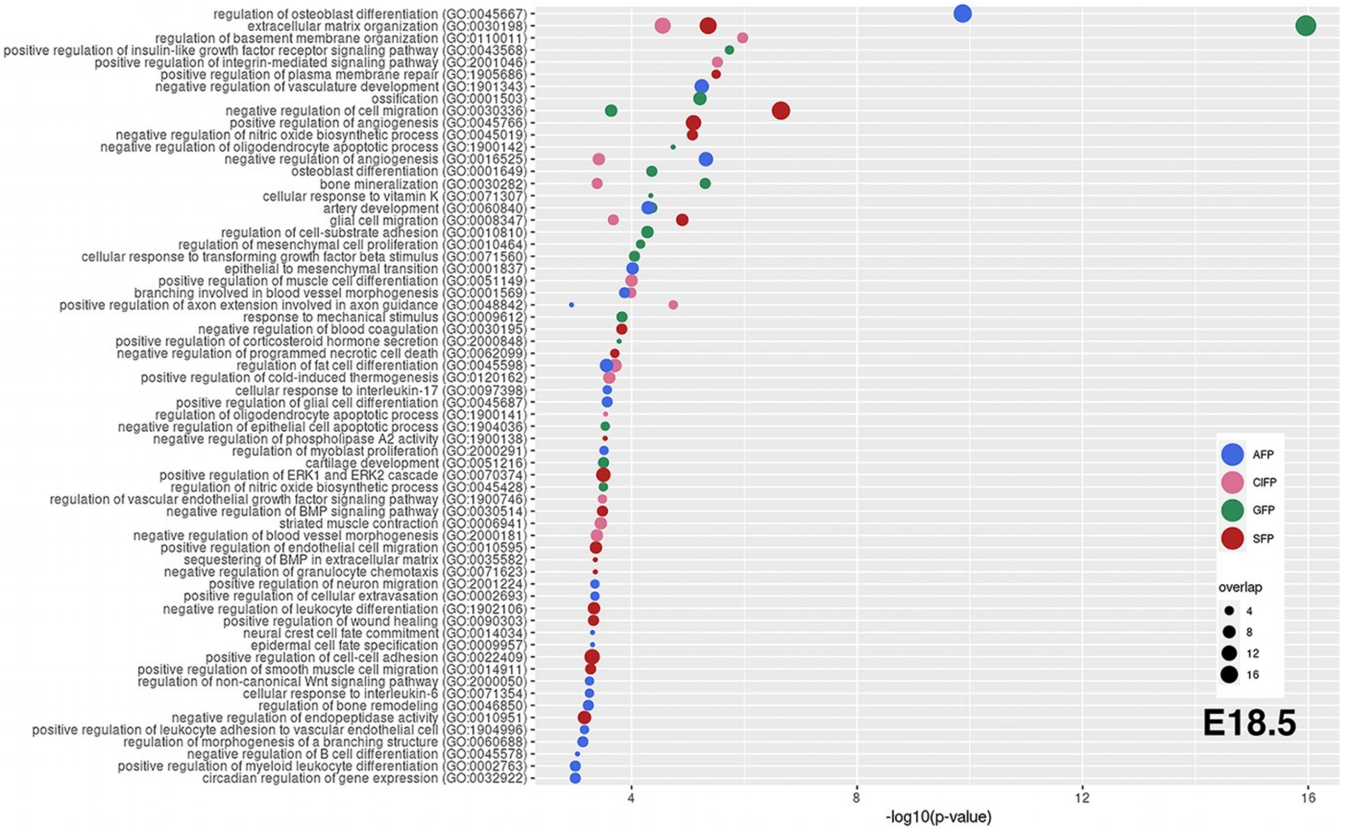


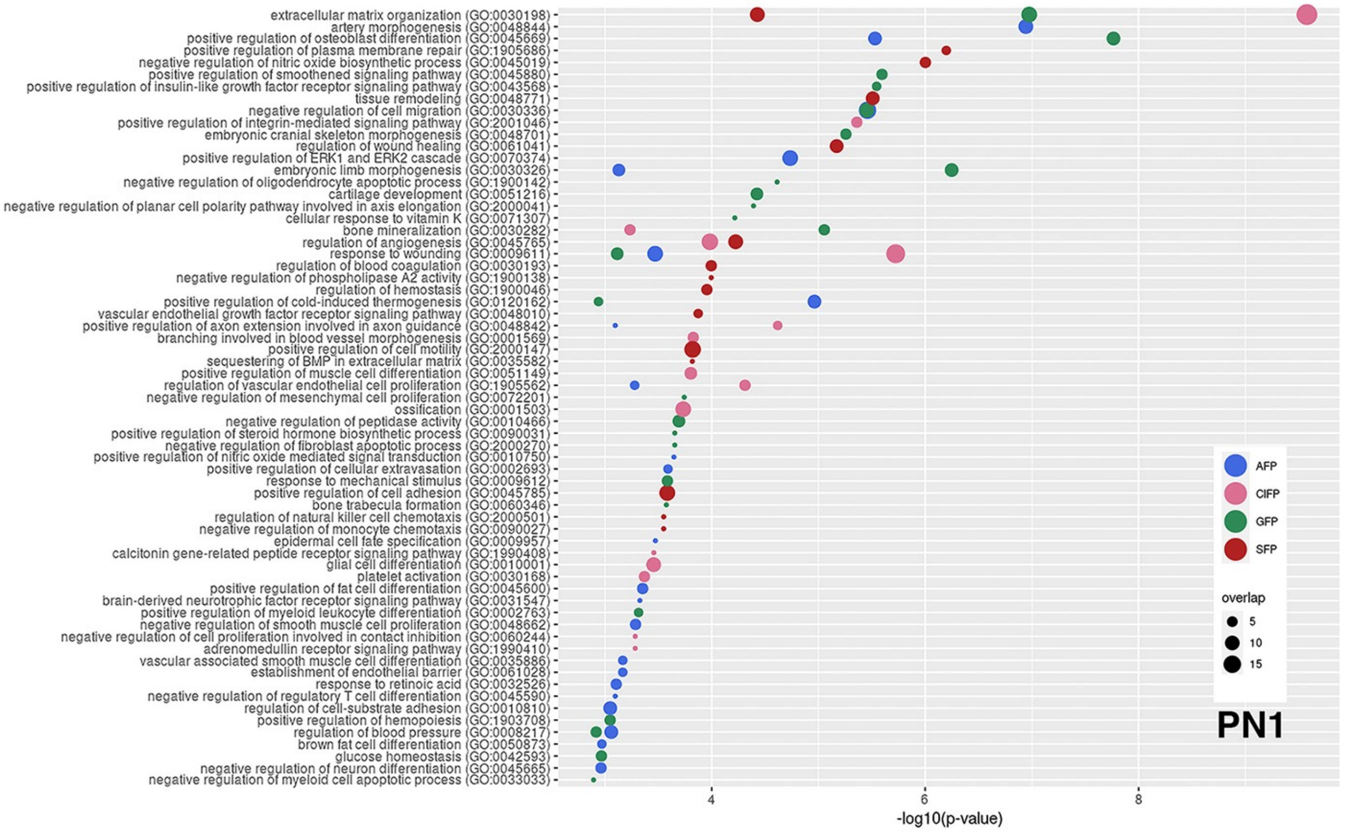


**Fig. S7. GO terms associated to the expression profiles of the GFP, SFP, GFP and CLFP fibroblastic clusters at E18.5 and PN1.**

### FIGURE S8

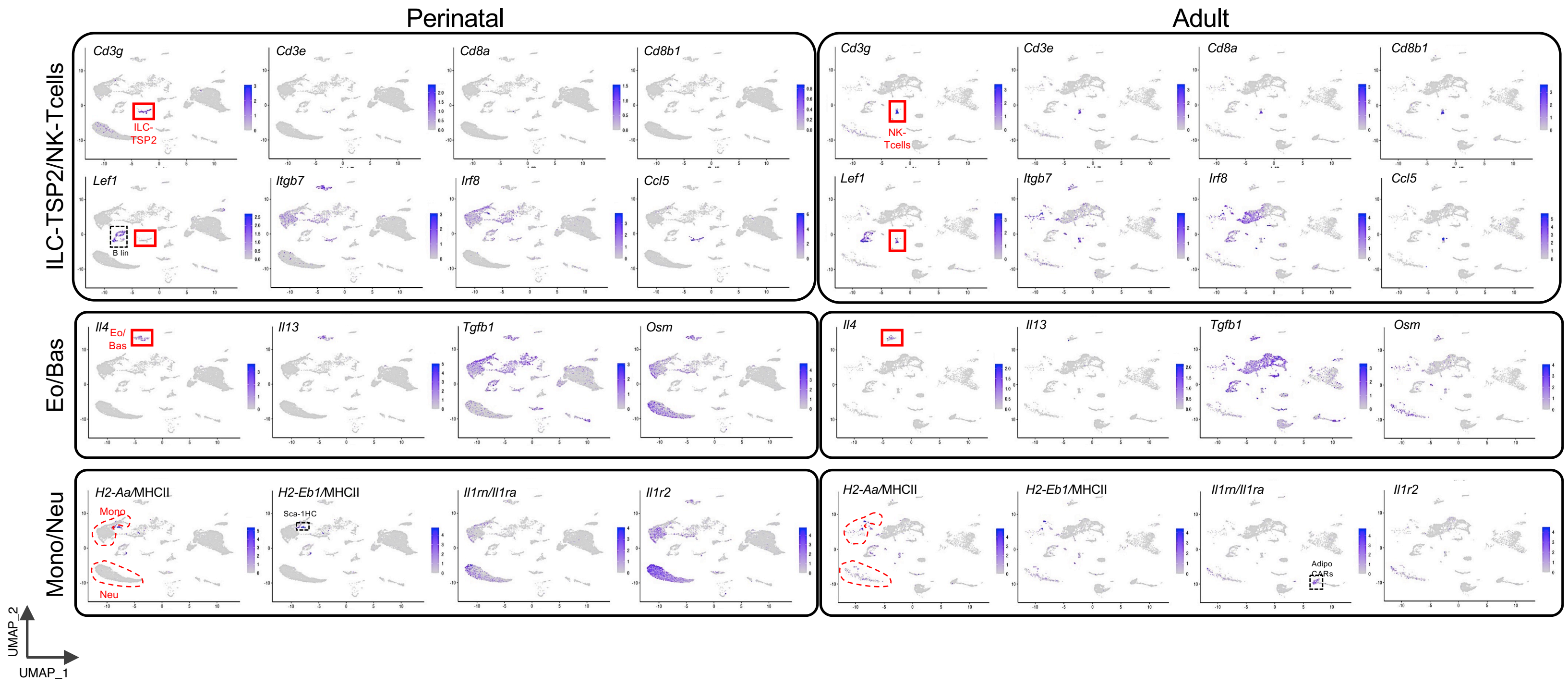


**Fig. S8. Expression differences between perinatal and adult hematopoietic clusters.** Upper panels correspond to ILC-TSP2 and NK- T cells (red box) in perinatal and adult bone, respectively. Note the absence of expression of *CD3e*, *Cd8a*, *Cd8b1* and *Lef1* in perinatal ILC-TSP2. B Lin (B lineage; black box) is labelled to show the expression of *Lef1* in both perinatal and adult stages. Middle panels show the expression of anti-inflammatory cytokine genes (*Il4*, *Il13* and *Tgfb1*) in Eosinophils and Basophils (Eo/Bas) clusters. *Osm* is expressed in Eo/Bas, Mono and Neu clusters at both stages. The bottom panels allow the comparison between Monocytes (Mono) and Neutrophils (Neu) clusters (encircled in red dashed lines). At perinatal stages, and in contrast to the adult, MHCII gene (*H2-Aa* and *H2-E*b1) expression is absent in these clusters, indicating their impairment in T-cell activation. MHCII genes are expressed in mature B cells of the B lin cluster and in Sca-1HC cells (black box). The IL1 decoy receptors-encoding genes *Il1rn/Il1ra* and *Il1r2* are expressed in these clusters at both stages, being *Il1r2* also expressed in adult AdipoCARs (black box in right bottom panels).

### FIGURE S9

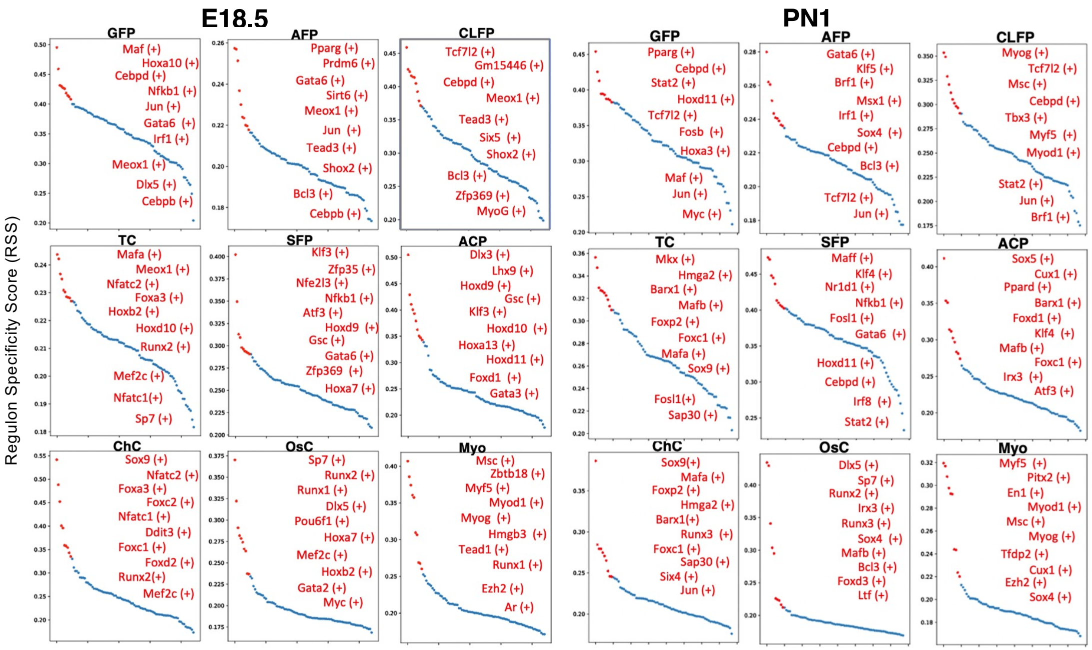


**Fig. S9. Transcriptional regulons operating in mesenchymal clusters at E18.5 and PN1 as analyzed by SCENIC.** On the left are shown the ten regulons with the highest RSS (regulon specificity score). The table on the right indicates specific regulons at E18.5 or PN1 which could be interesting for future studies. Represented in red are regulons among the top-ten; blue regulons are still significant (left panel). Notice that is some cases, there is a shift of paralog from E18.5 to PN1, as is the case of Runx1(+) and Runx3(+), or Klf3(+) and Klf4(+). Of note, the top regulons of TC and ACP at PN1 are Mkx (+) and Sox5 (+), respectively.

**Specific Regulons at E18.5 and PN1**

| Regulons | E18 Cluster | PN1 Cluster |
|---|---|---|
| Klf3 (+) | SFP ACP | |
| Klf4 (+) | | SFP ACP |
| Gsc (+) | SFP ACP OsC GFP | |
| Cux1 (+) | | ACP Myo |
| Runx1 (+) | OsC Myo | |
| Runx3 (+) | | OsC ChC |
| Nfatc1 (+) | ChC TC GFP | |
| Nfatc2 (+) | ChC | |
| Meox1 (+) | TC GFP AFP CLFP OsC | |
| Shox2 (+) | CLFP AFP GFP SFP | |
| Msx1 (+) | | AFP GFP SFP ACP |
| Stat2 (+) | | GFP CLFP AFP |
| Hoxa7 (+) | OsC SFP GFP Myo | |
| Tead1 (+) | Myo CLFP GFP | |
| Tead3 (+) | CLFP AFP GFP TC | |
| Mef2c (+) | OsC ChC TC | |
| Sox5 (+) | | ACP SFP ChC |
| Mkx (+) | | TC GFP ChC |
| Hoxa3 (+) | | GFP SFP TC AFP |
| Foxa3 (+) | ChC | |
| Irx3 (+) | | OsC ACP |
| Hoxa10(+) | GFP | |
| Barx1 (+) | | ACP ChC |
| Dlx3 (+) | ACP | |
| Foxd2 (+) | ChC GFP | |
| Foxd3 (+) | | OsC GFP CLFP TC |
| Ppard (+) | | ACP SFP |
| Foxc2 (+) | ChC OsC | |
| Foxp2 (+) | | ChC TC GFP |
| Nr1d1 (+) | | SFP ACP |

### FIGURE S10

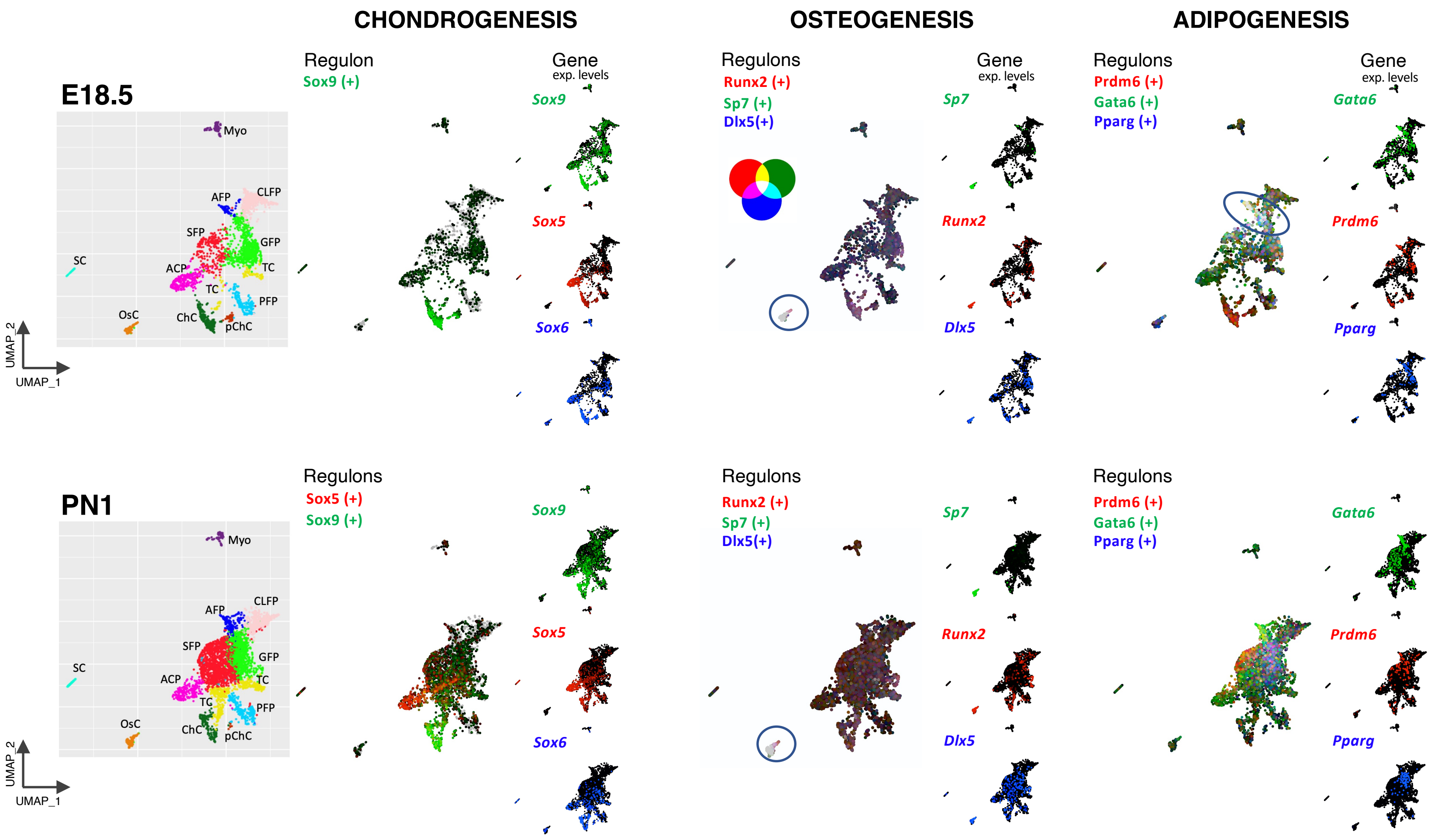


**Fig. S10. Chondrogenic, osteogenic and adipogenic regulons active in the perinatal endochondral bone.** UMAP projections of the mesenchymal clusters at E18.5 and PN1 are shown as a guide for cluster identification. SCENIC (default settings) displays up to three regulons simultaneously (the name of transcriptional factors and a (+) symbol) and the individual expression of their genes (in italic). For example, note that despite prominent *Sox5* expression at E18.5, its Sox5(+) regulon is not detected as active until PN1.

**Table S1. List of differentially expressed genes in AFP, CLFP, SFP and GFP clusters at E18.5 and PN1 (adjusted p-value ≤0.005) used for GO analysis.**
See supplementary file 'Table S1.xlsx'

**Table S2. Means and p-values of the CellPhoneDB analysis of all scRNAseq clusters at E18.5.** See supplementary file 'Table S2.xlsx'

**Table S3. Means and p-values of the CellPhoneDB analysis of all scRNAseq clusters at PN1.** See supplementary file 'Table S3.xlsx'

**Table S4. List of collagens, integrins and different ECM molecules within MC clusters at E18.5 and PN1 and provided by CellPhoneDB analyses**. Cluster-specific molecules or differences between intrauterine and extrauterine connectors are indicated in red. See supplementary file 'Table S4.xlsx'

**Table S5. Putative direct interactions between MC clusters at E18.5 as detected by CellPhoneDB (parameters: no secreted molecules; p-val<0,05; Log2mean>-1).** See supplementary file 'Table S5.xlsx'

**Table S6. Putative direct interactions between MC with EC or HC clusters at PN1 as detected by CellPhoneDB (parameters: no secreted molecules; p-val<0,05; Log2mean>-1).** See supplementary file 'Table S6.xlsx'
